# Structural basis for DNA double-strand break sensing by human MRE11-RAD50-NBS1 and its TRF2 complex

**DOI:** 10.1101/2025.03.14.643254

**Authors:** Yilan Fan, Filiz Kuybu, Hengjun Cui, Katja Lammens, Jia-Xuan Chen, Michael Kugler, Christophe Jung, Karl-Peter Hopfner

## Abstract

The MRE11-RAD50-NBS1 (MRN) complex is a central, multifunctional factor in the detection, signaling and nucleolytic processing of DNA double-strand breaks (DSBs). To clarify how human MRN binds generic and telomeric DNA ends and can separate DNA end sensing from nuclease activities, we determined cryo-electron microscopy structures of human MRN bound to DNA and to DNA and the telomere protection factor TRF2. MRN senses DSBs through a tight clamp-like sensing state with closed coiled-coil domains, but auto-inhibited MRE11 nuclease. NBS1 wraps around the MRE11 dimer, with NBS1’s ATM recruitment motif sequestered by binding to the regulatory RAD50 S site, necessitating an allosteric switch for ATM activation. At telomeric DNA, TRF2 blocks the second S site via the iDDR motif to prevent nuclease and ATM activation. Our results provide a structural framework for topological DNA sensing and separation of sensing, signaling and processing activities of mammalian MRN.

**Highlights:**

- Human MRN senses DNA ends with an autoinhibited nuclease
- NBS1’s C-terminus binds one RAD50 S site in the sensing state
- TRF2 binds MRN’s second S site at telomeres
- RAD50 and ATM compete for the NBS1 C-terminus

## Introduction

DNA double-strand breaks (DSBs) spontaneously arise from ionizing radiation, genotoxic chemicals, abortive topoisomerases or replication fork collapse, but are also programmed intermediates in meiotic recombination^1,2^ and immunoglobulin gene rearrangements^3-5^. Failure to properly repair DSBs can cause chromosomal aberrations, death or cancer^6,7^. Cells respond to DSBs through a cell cycle-regulated DNA damage response (DDR), primarily mediated by the activation of ATM, ATR, and DNA-PK kinases. This response leads to complex chromatin modifications^8,9^ and promotes either classic non-homologous end joining (c-NEHJ) or long-range resection followed by homologous recombination (HR)^10,11,12,13-16^. When both NHEJ and HR fail, cells can utilize error prone alternative end joining (alt-EJ) pathways^17^.

A central sensor of DSBs is the MRE11–RAD50–NBS1 (MRN) complex, known as MRX (Mre11-Rad50-Xrs2) in *S. cerevisiae.* MRN and its prokaryotic MR homologs are evolutionarily conserved ATP-dependent complexes with nuclease activity^18-24^. The core MR complex contains two copies of both the ATPase RAD50 and the endo-/exonuclease MRE11, assembled in an elongated structure. This structure consists of a “head” module which includes MRE11 and the ATP binding cassette type nucleotide binding domains (NBDs) of RAD50, and a “tail” module formed by the coiled-coil domains (CCDs) of RAD50^25-29^. NBS1^18^ is only found in eukaryotes, where it recruits ATM^30,31^ and the MRN nuclease cofactor CtIP^32-34^, among other factors. The CCDs of RAD50 are apically joined generating a ring-like protein structure that serves as an ATP-dependent gate for recognizing single DNA ends^35-39^. In bacterial MR, ATP binding opens the CCDs to load MR onto linear DNA, while ATP hydrolysis closes the CCDs to a rod-like state that repositions SbcD^MRE11^ from an autoinhibited “resting” to a “cutting” state^37,40^.

Following the detection of DSBs, MR/MRN nicks the 5’ strand in the vicinity of the break^2,41-48^, possibly followed by back-resection through its 3′→5′ exonuclease activity^24,49^, as well as cleavage of the exposed 3’ strand^45,50^. The nuclease activity removes covalent DNA-protein adducts^1,48^, opens and degrades hairpin structures, can generate ssDNA in alt-EJ pathways^51,52^ and activates the long-range resection machinery in HR^16^.

MRN/X has highly regulated, multifunctional roles in the DDR that besides sensing and nuclease activities also includes DNA tethering and ATM/Tel1 activation. Furthermore, MRN/X’s nuclease is highly regulated by trans-interacting factors that interact with a regulatory surface patch on RAD50, denoted “S site”. Structure prediction and functional studies suggest that the co-activator CtIP/Sae2 binds the S site to promote 5’ incision, while at telomeres, TRF2 (human) or Rif2 (yeast) bind the S site to prevent MRN-dependent activation of ATM and counteract CtIP/Sae2^53-61^.

To provide a structural framework for sensing of DNA ends by human MRN and its regulation, we determined cryo-electron microscopy (cryo-EM) structures of MR and MRN bound to DNA, as well as bound to DNA and TRF2. Our work identifies a “sensing state” that in this form is not observed in prokaryotic MR and can explain decoupling of MRN’s structural and nucleolytic functions at DSBs. Our structure shows that NBS1’s conserved C-terminal region, which carries the ATM recruitment motif, is sequestered at the RAD50 S site. This suggests an allosteric switch mechanism, where ATM and RAD50 compete for binding to NBS1’s C-terminus, regulating the formation of the MRN-ATM complex. We also provide a structural basis for the telomeric TRF2-MRN complex, showing that both RAD50 S sites are blocked through NBS1 on one side and the TRF2 iDDR motif on the other side.

## Results

### Structural basis for MRN’s interaction with DNA

Subunits of the MRN complex (Fig. 1A) were co-expressed in human or insect cells (Fig. S1A, see STAR Methods). The purified MRN complex showed the characteristic DNA-stimulated ATPase activity (Fig. 1B) and interaction architecture, determined by mass photometry and chemical cross-linking coupled with mass spectrometry (CX-MS) (Fig. S1B-D)^62^. To obtain DNA bound structures of MRN, we screened several different types of DNA and nucleotides. The highest resolution structures were obtained by incubating MR or MRN with 50 bp DNA and ATP at 35°C for 30 min, followed by addition of BeF_x_ (producing the ATP mimic ADP•BeF_x_ after ATP hydrolysis). Extensive processing resulted in 3D reconstructions with overall resolutions of 3.2 Å (MR) and 3.1 Å (MRN) (see STAR Methods and Figs. S2-S4). The maps resolve MR’s and MRN’s catalytic head bound to DNA and parts of the CCD module, with ADP•BeF_x_•Mg^2+^ bound at both active sites (Fig. 1C-F). We do not observe density for MRE11’s C-terminal extension (510-708), the N-terminal region of NBS1 (1-650), both of which are predicted to be unstructured or mobile, and the remainder of the RAD50 CCDs (∼245-∼1070). Atomic models were generated by rigid body fitting AlphaFold3 predictions^63^, manual fitting and automated refinement (see STAR Methods).

**Figure 1:**
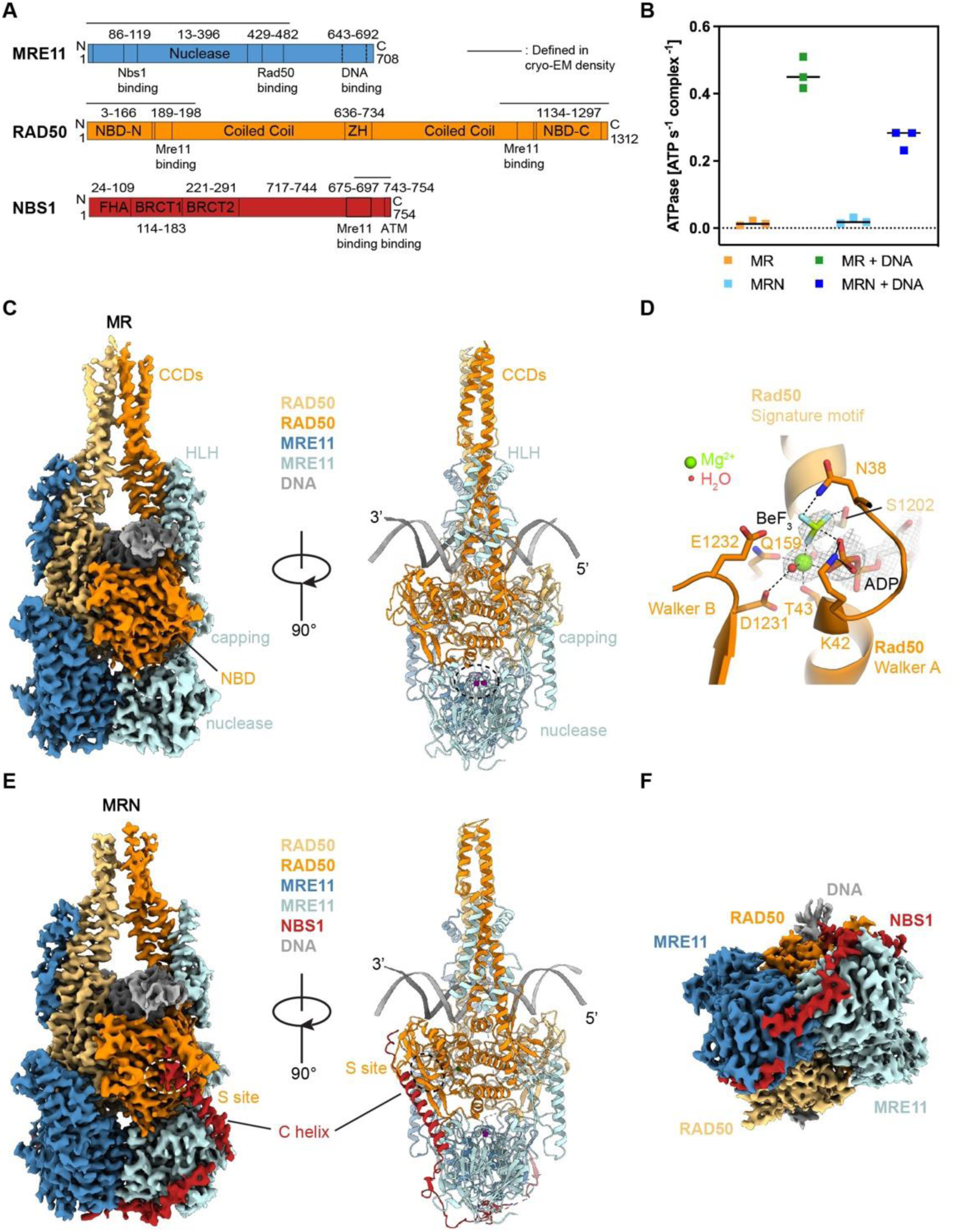
Overall structure of the human MR(N) complex bound to DNA. **A**, Domain arrangement of human MRN subunits. Domain boundaries are indicated with numbers and vertical black lines. ZH, zinc hook; FHA, forkhead-associated domain; BRCT, breast cancer C-terminal domain; NBS1 C-terminal helix, NBS1 C-helix. **B**, ATPase activity of MR(N) complex stimulated by DNA binding. **C**, Overview of the MR-DNA complex structure. High resolution cryo-EM map was displayed (left panel). MRE11 nuclease active site (buried) is highlighted in white/black dashed circle. **D**, Ribbon representation of the nucleotide-binding site of Rad50 highlighting the presence of BeF_3_ bound together with ADP. The density of ADP, BeF_3_, Mg^2+^ and water molecules from the structure of MR-TRF2^iDDR-Myb^-DNA complex (which has the best resolution) were contoured at 9 σ level. Black dashed lines represent electrostatic interactions or hydrogen bonds. **E**, Overview of the MRN-DNA complex structure. High resolution cryo-EM density was displayed (left panel). RAD50 S site is highlighted in white/black dashed circle. **F**, Top view of the MRN-DNA complex structure illustrated in **E** (left panel) obtained by a 90-degree rotation around the x axis.

hMR/MRN comprises two tightly engaged RAD50^NBD^s that bind into the concave side of the MRE11 dimer (Fig. 1C, E). Furthermore, MRE11 interacts with RAD50’s CCDs via a helix-loop-helix (HLH) domain, which is expanded to four helices in the human structure (Four-helix boundle). The nuclease active site of MRE11 harbors density consistent with two Mn^2+^ ions (present in the buffer, Fig. S4A) but is shielded from any DNA access by the proximity of RAD50^NBD^s.

DNA interacts with both RAD50^NBD^s in a two-fold symmetric manner, traversing the RAD50 dimer between the CCDs with a footprint of ∼20 bases (Fig. 1C, E, Fig. 2A, D-F), consistent with ATP dependent DNA binding (Fig. 2C). The 3’ strand, as viewed from a DSB, binds to the RAD50^NBD^ N-lobe along the top β-strand of the S site and to the subsequent α-helix, while the 5’ strand binds the N-lobe at a loop following the P-loop helix (Walker A motif) (Fig. 2D). Further contacts are observed at the RAD50^NBD^ C-lobe, where the 3’ strand binds to a helix connecting the Q-motif and CCD and a hairpin loop inserts R1179^RAD50^ into the minor groove (Fig. 2E, Fig. S4B). Latter interaction stabilizes a compression of the minor groove and an overall upward bending of DNA towards the periphery. MRE11 does not substantially contribute to DNA binding in this state, with only a single residue (K467^MRE11^) at the HLH element in sufficient proximity to the 3’ strand. DNA interactions are virtually identical in MR and in MRN complexes. However, the highly positively charged C-terminus of NBS1 (K_751_KRRR_754_) binds the DNA emerging from RAD50 and induces additional DNA contacts on one side of MRN (Fig. 2F).

**Figure 2:**
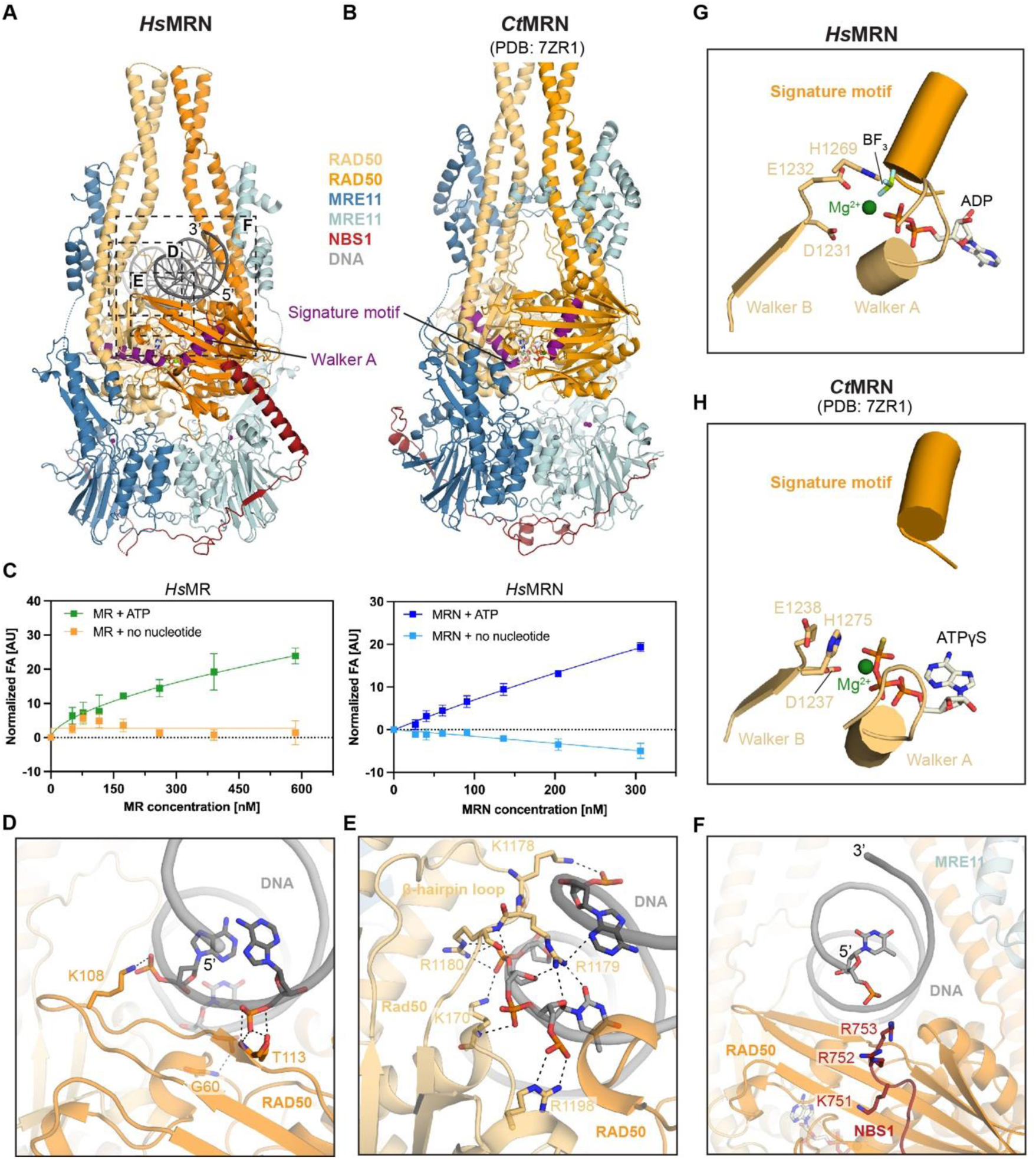
Structural basis of DNA binding by human MRN complex. **A and B,** Side by side comparison of the *Hs*MRN-DNA and *Ct*MRN (PDB: 7ZR1) complex structure. The α-helixes containing either the walker A or signature motifs in both structures were colored in purple for better visualization. *Ct*: *Chaetomium thermophilum*. **C**, Fluorescence anisotropy (FA)-based assay documenting the effect of ATP on MR/MRN binding towards telomeric repeat containing 64-mer DNA template. **D-F**, Detailed molecular contacts between DNA and the MRN complex. For simplicity, only the nucleotides of the DNA template involved in the interactions were displayed as sticks. Black dashed lines represent electrostatic interactions or hydrogen bonds. **G-H**, Structural comparison of nucleotide binding pockets in *Hs*MRN and *Ct*MRN (PDB: 7ZR1) structures.

Insertion of the RAD50^NBD^ hairpin loop (1168-1183) into the minor groove goes hand in hand with an inward tilting of both CCDs, which meet ∼20Å above the DNA (Figs. 2A, S4E). Together, RAD50^NBD^s and CCDs form a tight clamp around DNA that can accommodate only a single DNA duplex to pass consistent with CX-MS (Fig. S1C,D). In conjunction with cryo-EM studies of fungal (*Chaetomium thermophilum* Ct) MRN in the absence of DNA^64^, AFM studies on human MR/MRN with DNA^38^ and crystallographic analysis of the zinc-hook region^36^, the structure argues that CCDs may adopt a rod state along their length. Since we do not observe DNA contacts that directly probe the chemical nature of a DNA end, our structural results argue for a topological sensing of linear DNA through ATP dependent loading coupled with ring←→rod transition as observed for the bacterial MR^SbcCD37,40^. Here, restricting DNA passage through the CCD rod to a single DNA duplex ensures that the stable clamp can only form after loading onto linear DNA, but not internal or circular DNA. In latter case, at least two dsDNA segments would traverse the CCDs, preventing rod formation through steric hindrance.

### A “sensing state” decouples resting and cutting conformations

We observe density for ADP•Mg^2+^, with the better resolved structures also showing density for BeF_x_ (Fig. 1D). ADP•Mg^2+^•BeF_x_ moieties are bound to Walker A and B motifs of one RAD50^NBD^, and to the signature motif helix on the other RAD50^NBD^ (Fig. 2G). H1269^RAD50^ is positioned to act as γ-phosphate sensor. The active site conformation is conserved in all MR, MRN and MRN-TRF2 complex structures, suggesting that NBS1 and TRF2 do not allosterically modulate RAD50’s ATPase site. The RAD50 conformation in the human MR/MRN-DNA complexes is very similar to that of the crystal structure of fungal RAD50^NBD^ bound to DNA and ATPγS, except for the CCD angle^65^, which points outwards in the latter structure (Fig. S5). However, it is quite distinct from that of fungal MRN bound to ATPγS in the absence of DNA^64^ (Fig. 2B). In that structure, the CCDs were further inward positioned while the RAD50^NBD^ s were in an ATPase inactive state. This is shown by the strongly divergent signature motif and Walker A motif helices and the lack of direct interactions of the signature motif to the nucleotide phosphates in *Ct* MRN (Fig. 2H). The conformational differences suggest that DNA binding to RAD50^NBD^ leads to rearrangement of both lobes to properly align the ATPase motifs for catalysis as basis for DNA stimulated ATPase activity.

In several datasets, we observe 2D classes with open CCDs, in particular in the presence of ATPγS, more rarely in the presence of ATP+BeF_x_ (Fig. S6 A-G). In latter case, the Mre11 dimer is also sometimes tilted with respect to Rad50^NBD^s (Fig. S6C). We could not perform a convincing 3D reconstruction using these classes due to high orientational bias and flexibility. Evidently, ATP binding to MR/MRN in the absence of DNA can widen the CCDs to enable DNA loading, similar to the resting state of SbcCD^37^. DNA binding prompts closure of the CCDs and induces a conformation with aligned ATPase motifs but still enables some Mre11 movements. However, in contrast to bacterial MR, closing of the CCDs to a rod does not yet reposition the MRE11 nuclease from the auto-inhibited state to the cutting state. We therefore refer to this new state observed in human MRN as the “sensing” state.

### NBS1 tethers MRE11 to the RAD50 S site via the ATM recruitment motif

Our structure resolves a single NBS1 polypeptide (651-754), which wraps around the MRE11 dimer with the highly conserved K_683_NFKKFKK_690_ motif situated across the two-fold symmetry axis (Fig. 1F). NBS1 does not induce significant changes into the DNA bound MR structure, compared to MR alone (Fig. S10D).

Intriguingly, the very C-terminal region of NBS1 (718-745) forms a continuous helix (“C helix”) that bridges one MRE11 catalytic domain to one RAD50^NBD^ in the sensing state (Fig. 3A). The first part of the helix binds MRE11 via a hydrophobic interface between L719^NBS1^, L732^NBS1^, W722^NBS1^ and F233^MRE11^ (Fig. 3C). The second part of the C helix is bound to the N-lobe β-sheet at the RAD50 S site (Fig. 3D). Here F744^NBS1^ binds into an S site pocket lined by K6^RAD50^, S8^RAD50^, I24^RAD50^, R83^RAD50^ and Q85^RAD50^. Further notable interactions are electrostatic interactions between K6^RAD50^ and D741^NBS1^, and between R83^RAD50^ with the dipole charge of the NBS1’s C helix.

**Figure 3:**
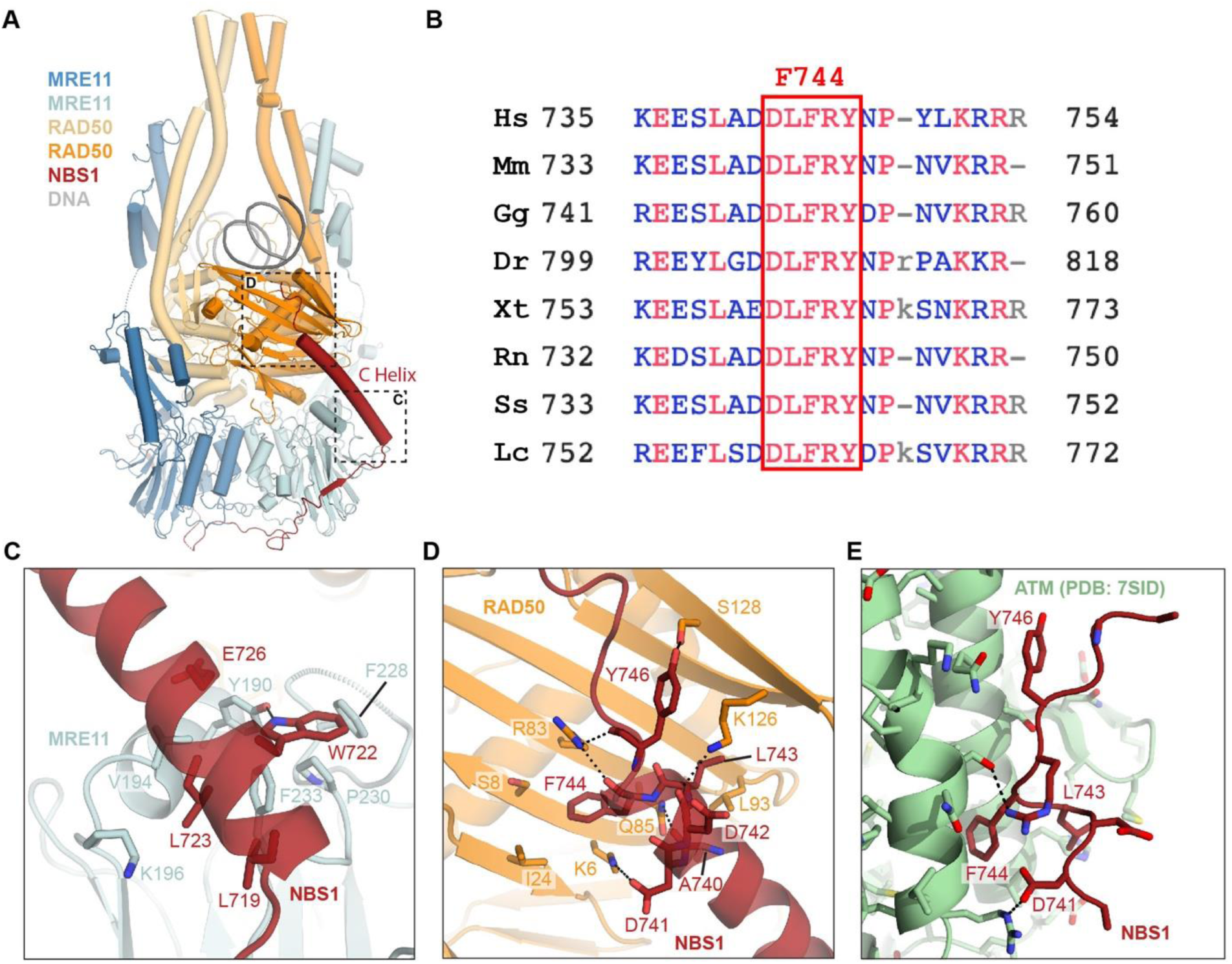
NBS1 interactions with the MR complex mediated by the C-helix. **A,** Overview of the MRN-DNA complex structure in ribbon model. **B**, NBS1 sequence alignment (*Hs*: *Homo sapiens, Mm*: *Mus musculus, Gg*: *Gallus gallus, Dr*: *Danio rerio, Xt*: *Xenopus tropicalis, Rn*: *Rattus norvegicus, Ss*: *Sus scrofa, Lc*: *Latimeria chalumnae*) highlighting the conservation of F744 (numbered according to *Hs*NBS1). **C and D**, Close-up views of the interactions formed between NBS1 C-helix and MRE11 or RAD50, respectively. Only interacting residues were displayed as sticks here for simplicity, and the black dashed lines represent electrostatic interactions or hydrogen bonds. **E**, Associations of NBS1 C-helix with ATM (PDB: 7SID) illustrated in a similar view to **D** for the purpose of side-by-side comparison.

During 3D classification, we identified two separable conformational states in MR and MRN, one with the MRE11 dimer symmetrically attached to both RAD50^NBD^ s, and one where the MRE11 catalytic domain dimer detaches as a rigid body from one RAD50^NBD^ (Fig. S6H). In the case of MRN, the detachment is only observed at the RAD50^NBD^ not bound to NBS1. These observations suggest a stabilizing effect of the NBS1 C helix-S site interaction on MRN catalytic head dynamics. They also indicate an asymmetry of detachment and putative path for a conformational transition to a cutting state.

Since the C helix contains an important ATM recruitment motif^30,31,66^ we compared S site binding of the C helix to binding of the NBS1 C-terminal peptide to the ATM dimer (PDB accession code: 7SID)^67^ (Fig. 3D, E). Both interactions are mediated by a highly conserved DLFRY sequence in NBS1 (Fig. 3B). F744 at the center of the motif provides a hydrophobic anchor in both MRN and ATM/NBS1 structures, binding to pockets in either the RAD50 S site or in the regulatory N-terminal HEAT repeat region of ATM. Evidently, RAD50 in the sensing state and ATM would need to compete for the C helix and ATM recruitment by MRN necessitates detachment of the C helix of NBS1 from the S site. Altogether, this suggests an allosteric transition in MRN and detachment of NBS1 from the RAD50 in the DNA sensing state to a signaling state in complex with ATM.

### Structures of the TRF2-MR and TRF2-MRN complex bound to DNA

To provide a structural basis for inhibition of ATM signaling and DNA cleavage at telomeric DNA, we determined cryo-EM structures of MR and MRN bound to the shelterin^68,69^ component TRF2 and DNA containing TTAGGG motifs (Fig. 4 A-D and S7-S9). TRF2 interacts with the S-site via the iDDR motif based on structure prediction and mutagenesis, but a comprehensive architecture including the conformational state of MR or MRN remains open ^53,61^. TRF2 contains a telomeric repeat factor homology (TRFH) dimerization domain, an iDDR motif and Myb domain (490-542) that binds double-stranded telomeric TTAGGG repeats. Adding the TRF2^iDDR^ (438-485) to our cryo-EM sample preparation procedure led only to free MRN-DNA and MR-DNA structures without additional density for TRF2^iDDR^. We also did not observe an interaction of TRF2^iDDR^ with MRN in gel filtration, although more sensitive yeast two-hybrid assays can detect RAD50 TRF2 interactions^61^. We reasoned that by including the Myb domain, we can stabilize the interaction through avidity effects, essentially recapitulating what happens at telomeric DNA. Using a TRF2 construct harboring both the iDDR and Myb domains (TRF2^iDDR-Myb^), as well as full length TRF2, we could determine high resolution structures of MR-TRF2^iDDR-Myb^ -DNA, MRN -TRF2^iDDR-Myb^ -DNA and MRN -TRF2-DNA (Fig. S7-S9).

**Figure 4:**
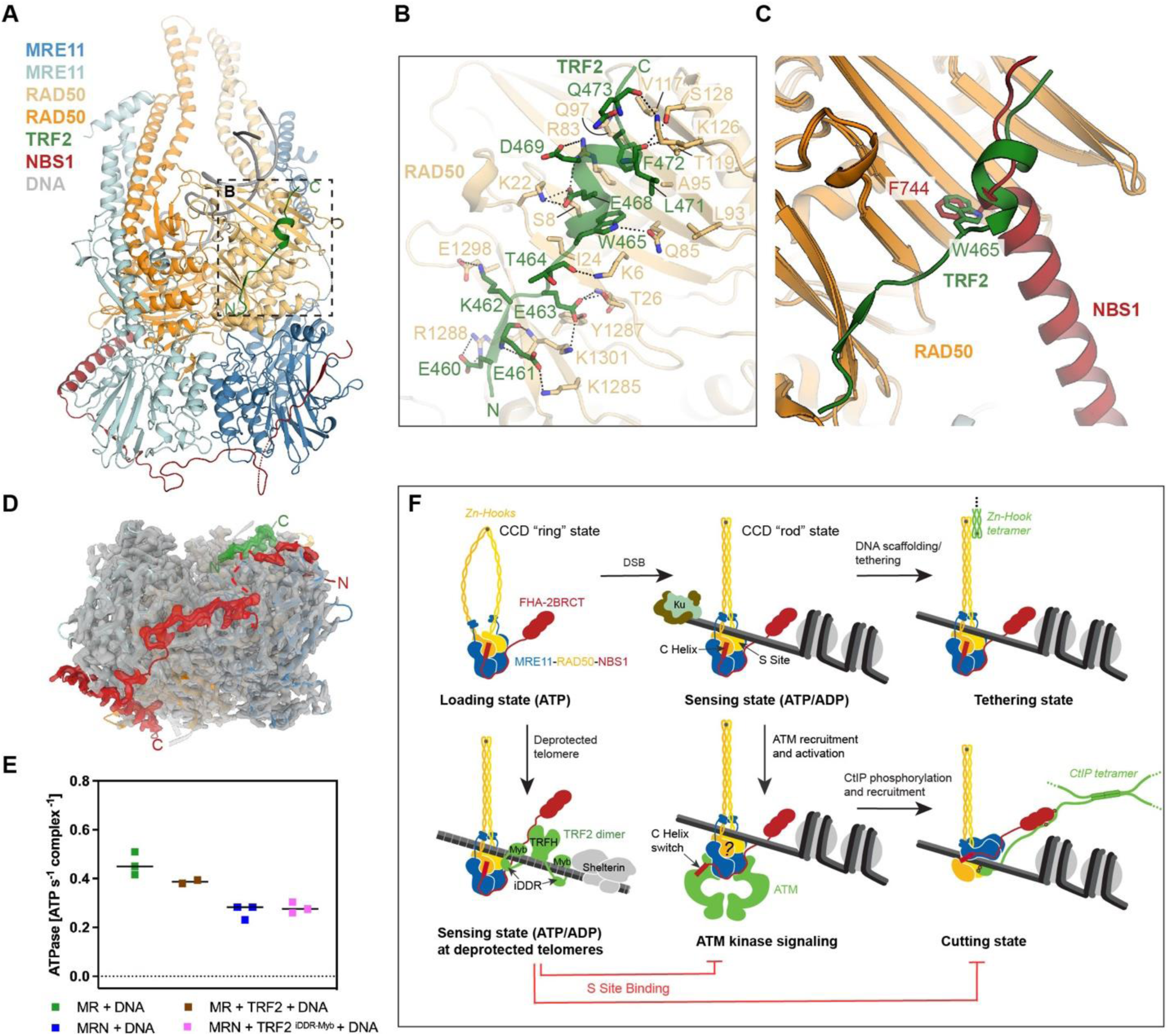
Structural basis of TRF2^iDDR-Myb^ binding to the human MRN complex. **A,** Overview of the MRN-TRF2^iDDR-Myb^-DNA complex structure. **B**, Zoomed-in view of the molecular interactions between TRF2^iDDR^ and RAD50. The residues involved in the associations were illustrated as sticks, black dashed lines represent electrostatic interactions or hydrogen bonds. C, Comparison of binding sites to RAD50 between NBS1 C-helix and TRF2 centering the evolutionary conserved hydrophobic residue F744 (*Hs*NBS1) or W465 (*Hs*TRF2). **D**, Top view of the MRN-TRF2^iDDR-Myb^-DNA complex structure emphasizing the asymmetric binding property of NBS1 and TRF2. The high resolution cryo-EM density is displayed in grey except for that of the resolved parts of NBS1 and TRF2, which were colored in red and green, respectively. **E**, Comparison of ATPase activities by MR/MRN complex in the presence or absence of TRF2. **F**, A proposed working model of the MRN complex initiated by DSB in chromosome.

The reconstructions show well-resolved density for TRF2 459-473, essentially comprising the iDDR region (Fig. 4A, B). Myb and TRFH domains are not resolved, indicating a flexible and structurally independent arrangement that averages out in 3D reconstructions. The core of the iDDR S site interaction is a short helical element that forms a hydrophobic-aromatic anchor through W465^TRF2^, L471^TRF2^ and F472^TRF2^ with a small hydrophobic patch on RAD50 (L93, A95, V117). Contribution to the core interaction is an acidic cluster (E_467_EDE_470_^TRF2^) that forms electrostatic interactions with K6^RAD50^, K22^RAD50^, R83^RAD50^ and K126^RAD50^. The region C-terminal to this core interaction points towards the DNA section that is exiting MR/MRN, consistent with the Myb domain binding to DNA in the wider vicinity of MRN. We observe density up to Q475^TRF2^. The subsequent 15 amino acid linker and Myb domain C-terminal to the RAD50 binding region are not resolved. The N-terminal part of iDDR reaches down all the way to the MRE11 dimer but we do not observe density that would indicate stable interactions with MRE11.

Intriguingly, in the case of MR, both RAD50^NBDs^ are occupied by iDDR, while in the case of MRN, density for iDDR was only observed at one S site, whereby the other one is still bound by the NBS1 C helix (Fig. 4D). This is due to steric competition, and not allosteric regulation, with NBS1 evidently outcompeting TRF2. While the binding sites of NBS1 and TRF2 strongly overlap (Fig. 4C), with hydrophobic anchor residues (F744^MRE11^ or W465^TRF2^) occupying the same S site pocket, “MR” and “MRN” parts of the TRF2 complexes are practically identical to the corresponding structures obtained in the absence of the TRF2/TRF2^iDDR-Myb^. The similarity includes also nucleotide state and active site geometry (Fig. S10D). Consistently, we neither observe a stimulation nor a reduction of MRN’s ATPase by TRF2/TRF2^iDDR^ (Fig. 4E). Along the same lines, in TRF2/TRF2^iDDR-Myb^ complexes, we also observe the mobility of MRE11 dimer, which detaches from the TRF2 bound but not the NBS1 bound RAD50^NBD^ (Fig. S6H).

The particular “rigid-body”-like recognition of the sensing state by TRF2 suggests that TRF2 does not directly provoke MRN release from DNA. To test this, we performed DNA binding assays in the presence of ATP. TRF2 bound to DNA with an affinity of 121 nM (Fig. S10A). Titrating MRN in the presence or absence of a saturating concentration of TRF2 both resulted in MRN binding to DNA, suggesting that even saturating TRF2 concentrations do not block MRN. However, while MRN binding does not lead to a plateau in our experimental conditions, suggesting multiple binding or clustering events, we obtain a proper binding curve in the presence of TRF2 (Fig. S10B), suggesting a more defined, single binding event and complex.

Taken together, TRF2 binds MRN in a 1:1 complex and prevents ATM and CtIP associated activities by saturating and blocking the free S site, without impacting the sensing state itself.

## Discussion

Here we provide cryo-EM structures of the human MRE11-RAD50-NBS1 (MRN) DSB sensor and repair factor bound to DNA and TRF2, altogether providing a structural framework for DNA end sensing by MR/MRN at generic DNA double-strand breaks or at de-protected telomeres. The structures are generally consistent with the “topological” sensing of DSBs as proposed for the bacterial MR^SbcCD^ homolog. DSBs are detected via ATP-dependent ring←→rod transitions of the CCDs that allow loading of MR/MRN onto linear DNA, but not internal/circular DNA. ATP-dependent widening of the RAD50 CCDs enables DNA binding to the RAD50^NBD^s. In case of binding near or at an DNA end, a single DNA duplex would pass through the CCD ring, which can subsequently close to a rod with a channel for a single DNA duplex as observed in the structures. If circular/internal DNA is bound, DNA would pass through the CCD ring twice. The dual passage could prevent rod formation and formation of a stable clamp, promoting DNA release. This mechanism could explain why MRN can load onto DNA ends sterically blocked by proteins and cleaves DNA at some distance from the DNA end^45,70,71^.

A striking contrast of human MRN’s clamp around DNA compared to bacterial MR is the location of the MRE11 nuclease. In bacterial MR, formation of the CCD rod coincides with a movement of the MRE11 dimer to the cutting state conformation, whereby DNA end sensing and cleavage are functionally coupled^37,40^. Evidently, in human MR/MRN, the MRE11 dimer remains in the auto-inhibited state with nuclease active sites blocked by RAD50, although the CCDs form a related rod like conformation. We refer to this new state of MRN/MR’s as “sensing” state to distinguish the DNA bound state from the resting (ATP bound but no DNA) and cutting states. Importantly, it solves the conundrum how mammalian MRN on one hand can sense DNA ends through CCD ring→rod transitions yet avoid automatic nuclease activation like in the bacterial system. Furthermore, the sensing state would be very well suited to adopt non-nuclease functions at DNA breaks such as the tethering or signaling functions of the MRN complex (Fig. 4F).

In the sensing state, the active sites of the best resolved structures contain density for ADP•BeF_x_. A recently published yeast MR-DNA complex bound by ATPγS and carrying a Walker B DE->DQ mutant has a similar “clamp” conformation with closed CCDs, while bacterial MR adopted a RAD50 clamp state only after ATP hydrolysis and simultaneous MRE11 relocation (preprint DOI: 10.21203/rs.3.rs-5390974/v1). However, in the presence of ATPγS, we only observed classes with open CCDs at this point, indicating that ATP analogs only inconsistently recapitulate ATP and somewhat differ from species to species. Based on the variability analysis and 2D classes, we note that apparently the MRE11 dimer is not fully fixed in the sensing state but moves with respect to the RAD50 dimer. It is plausible that ATP hydrolysis or repeated ATP cycles induce dynamics that enable ATM or CtIP to fully induce structural conformations that act in ATM activation and nuclease reactions. However, the MRE11 dimer itself appears to be stable even in the absence of NBS1, thus further studies are needed to reveal how the cutting state is induced.

A single NBS1 subunit wraps around the MRE11 dimer, similar to what has been observed for fungal MRN^64^. The unexpected interaction of NBS1’s C helix with RAD50, which was not observed in the fungal complex, explains a previously observed direct interaction between RAD50 and NBS1 in the absence of MRE11^72^. The interaction is intriguing, as it involves two important regulatory motifs of MRN that mediate interactions with other factors (ATM, TRF2, CtIP). The S site on RAD50 and the ATM recruitment motif on NBS1 bind to each other, shielding themselves from direct interaction with other factors. It restricts interactions with TRF2 to one side of the complex by outcompeting TRF2 from the occupied S site. AlphaFold 2/3 predictions of the interaction of iDDR with the S site were relatively accurate and extensive mutational studies have validated this interaction. The observed structure can also explain effects of early studies of RAD50 S site mutants in mice, where mice carrying the *Rad50*^K22M^ allele along null were viable, but exhibited telomere failures and ATM dependent apoptosis, while other MRN associated activities were less affected^73,74^. K22^RAD50^ forms a specific salt bridge with E468^TRF2^, but not with NBS1. According to AlphaFold3 predictions, K22^RAD50^ also does form a direct salt-bridge with CtIP, which may explain these observations with a preferential or stronger destabilization of the interaction with TRF2^iDDR^.

Interestingly, TRF2 binds the sensing state without inducing significant conformational changes in either CCDs, RAD50^NBD^s or MRE11. We also do not observe a stimulation of MRN’s ATPase activity by TRF2 (Fig. 4E). A 2-3 fold stimulation of *S. cerevisiae* MR ATPase by Rif2 was observed *in vitro*, so there could be species-specific differences^55^. In any case, our observations argue against a model, where TRF2’s function is to remove MRN from telomeres. Rather, it sequesters the sensing state at unprotected telomeres and might even stabilize it, consistent with TRF2-dependent co-localization of MR/MRN at telomeres^54^, where MRN could adopt structural functions. Additional cell cycle-regulated interactions of TRF2’s TRFH domain with NBS1 Y_429_QLSP_433_^75^, prevent access of telomere specific exonuclease Apollo in G1. These additional interactions, consistent with a 1:1 MRN-TRF2 complex, would strengthen the interaction between MRN and TRF2 and suggest that a segregated, nuclease-inhibited sensing state could help protect or assist with structural functions in de-protected G1 telomeres (Fig. 4F).

Our structural results reveal that due to steric competition, NBS1’s C helix needs to dissociate from the S-site, indicating the necessity of a structural switch upon formation of the MRN-ATM complex (Fig. 4F). An NBS1 variant (1-726, NBS1^ΔC^) that lacks the RAD50 interacting C-helix resulted in a hypomorphic phenotype with most MRN functions intact, but improper phosphorylation of a subset of ATM targets such as SMC1^66^, while the deletion also mitigated the effect of the RAD50S mutation. Since the C-terminal motif is not sufficient for activation of MRN/Tel1 by MRN/MRX and other parts of the complexes are needed as well, it is plausible that detachment of the NBS1 C helix on one hand leads to a more stable ATM interaction at DNA ends and phosphorylation of the full set of targets, but also frees up a binding site or otherwise enables formation of structural state in MRN that binds ATM for full activation. Along the same line, the C helix would need to detach from the S site to enable MRE11 to relocate to the other side of DNA to form the cutting state. However, since deletion of parts of the C helix did not lead to increased genome instability in mice^66^, it might not play a critical role in the cutting state, once it is formed.

In summary, we identify a DSB sensing state of MRN and MRN-TRF2 that argues for the DNA end chemistry insensitive, topological detection of DSBs, whereby the S site of Rad50 emerges as structural hub for both intrinsic and extrinsic regulators (Fig. 4F). The structural analysis solves the conundrum how DNA end detection and nuclease functionalities are separated in eukaryotes and mammals and provide a framework to understand subsequent regulatory steps.

### Limitations of the study

Our structures do not yet explain the precise roles of ATP hydrolysis in DNA end processing, nuclease activities, and what structural transitions happens to form the cutting state and signaling states of MRN. Further work is needed to understand a possible mechanistic function of a TRF2 bound sensing state at de-protected telomeres.

## STAR ⋆ METHODS

Detailed methods are provided in the full version of this manuscript and include the following:

- KEY RESOURCES TABLE
- RESOURCE AVAILABILITY

- Lead contact
- Materials availability
- Data availability
- EXPERIMENTAL MODEL AND SUBJECT DETAILS
- Experimental models: Organisms as source for materials used in experiments
- METHOD DETAILS

- Expression and purification of MR and MRN complexes
- Expression and purification TRF2 complexes
- DNA Substrates
- Fluorescence anisotropy DNA binding assays
- NADH-coupled ATPase assay
- Cryo-EM grid preparation
- Cryo-EM data acquisition collection
- Cryo-EM image processing
- Model building
- Crosslinking mass spectrometry
- In gel digestion
- Liquid chromatography tandem mass spectrometry
- Mass spectrometry data analysis
- QUANTIFICATION AND STATISTICAL ANALYSIS

## SUPPLEMENTAL INFORMATION

## ACKNOWLEDGEMENTS

We thank the Cryo-EM Core Facility of the Gene Center, Department of Biochemistry, LMU, Munich for technical support during EM data collection and all members of the Hopfner lab for helpful discussions. We are grateful to Olga Fettscher, Brigitte Kessler, Halina Kurzyca-Börncke, Manuela Moldt and Alexandra Scheele for excellent technical assistance. Y.F and F.K acknowledge support from the International Max-Planck Research School for Molecules of Life. Research was funded by the Deutsche Forschungsgemeinschaft (CRC1361, Gottfried Wilhelm Leibniz-Prize, HO2489/11-1 to K.P.H.). Funding of CRC1361 supported the Exploris 480 system (J.X.C).

## AUTHOR CONTRIBUTIONS

Y.F and F.K conducted all experiments (MR and MRN purification, cryo-EM sample preparation, *in vitro* biochemistry assays, chemical crosslinking) and together with M.K and K.L collected cryo EM data. Y.F, F.K, H.C, and K.P.H processed EM-data, with advice from M.K.. H.C purified TRF2 variants and carried out structural model building, refinement and deposition. J.X.C performed XL mass spectrometry analysis. C.J. performed fluorescence anisotropy data collection. Y.F, F.K, H.C, K.P.H, and K.L analyzed and interpreted data. K.P.H designed and supervised the overall project and provided funding. Y.F, F.K, H.C and K.P.H wrote the manuscript with input from all authors.

## RESOURCE AVAILABILITY

### Lead contact

Further information and requests for resources and reagents will be fulfilled by the lead contact, Karl-Peter Hopfner.

### Materials availability

Plasmids and other materials that were generated in this study are available without restrictions from the lead contact.

### Data availability

Cryo-EM data generated in this study were deposited as coordinate files in the Protein Data Bank (https://www.rcsb.org) and reconstructions in the Electron Microscopy Data Bank (https://www.ebi.ac.uk/pdbe/emdb/) under the following accession codes for each structure:

MR-DNA: EMDB: 52959 (3.21 Å) and PDB: 9Q9H.

MRN-DNA complex: EMDB: 52960 (3.08 Å) and PDB: 9Q9I.

MR-TRF2^iDDR-Myb^-DNA complex: EMDB: 52962 (2.7 Å) and PDB: 9Q9K.

MRN-TRF2^iDDR-Myb^-DNA complex: EMDB: 52961 (3.01 Å) and PDB: 9Q9J.

MRN-TRF2-DNA complex: EMDB: 52964 (2.97 Å) and PDB: 9Q9M. Additional information required for the analysis of reported data is available upon request from the lead contact.

## STAR * METHODS

### KEY RESOURCES TABLE

#### Chemicals and Reagents

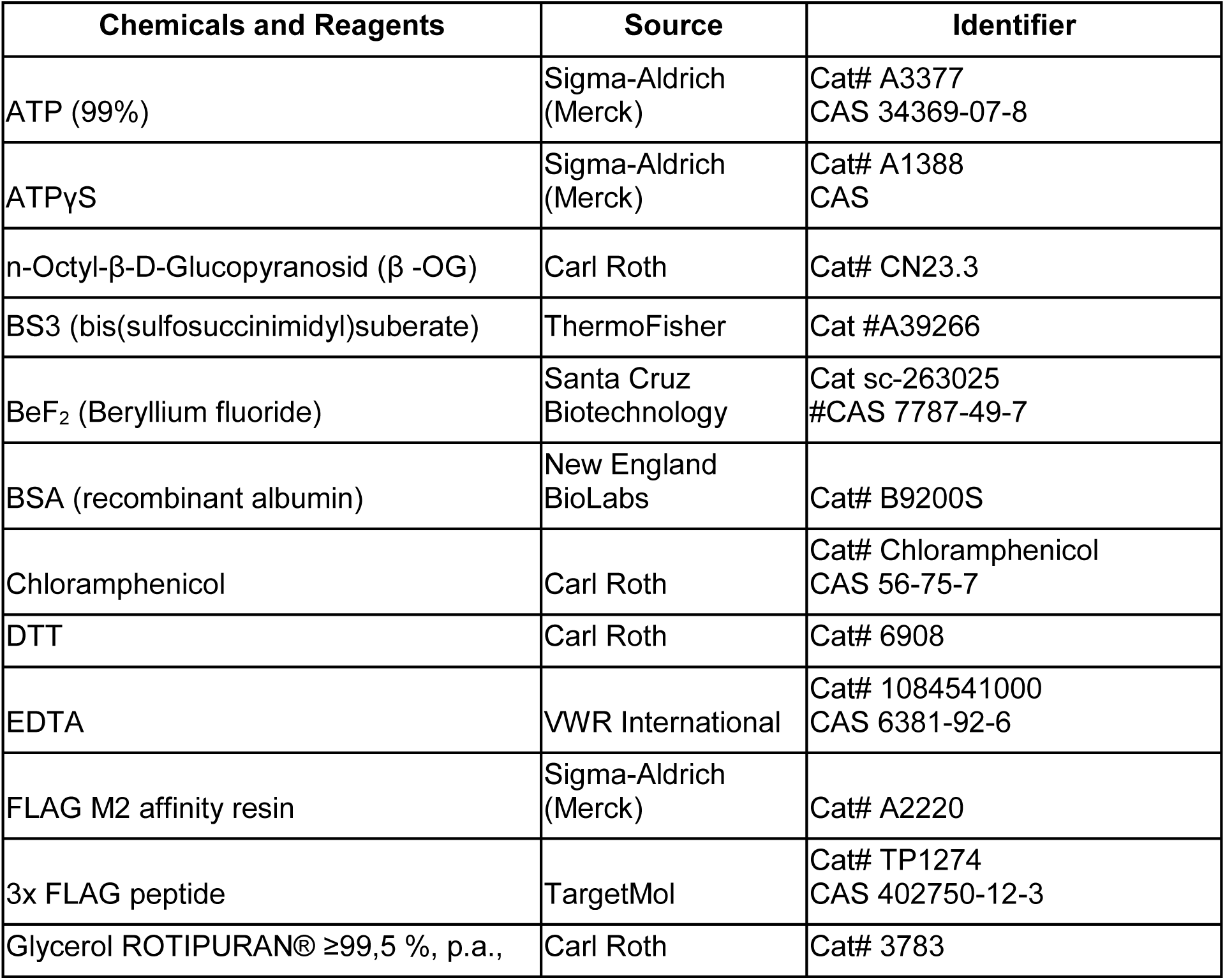
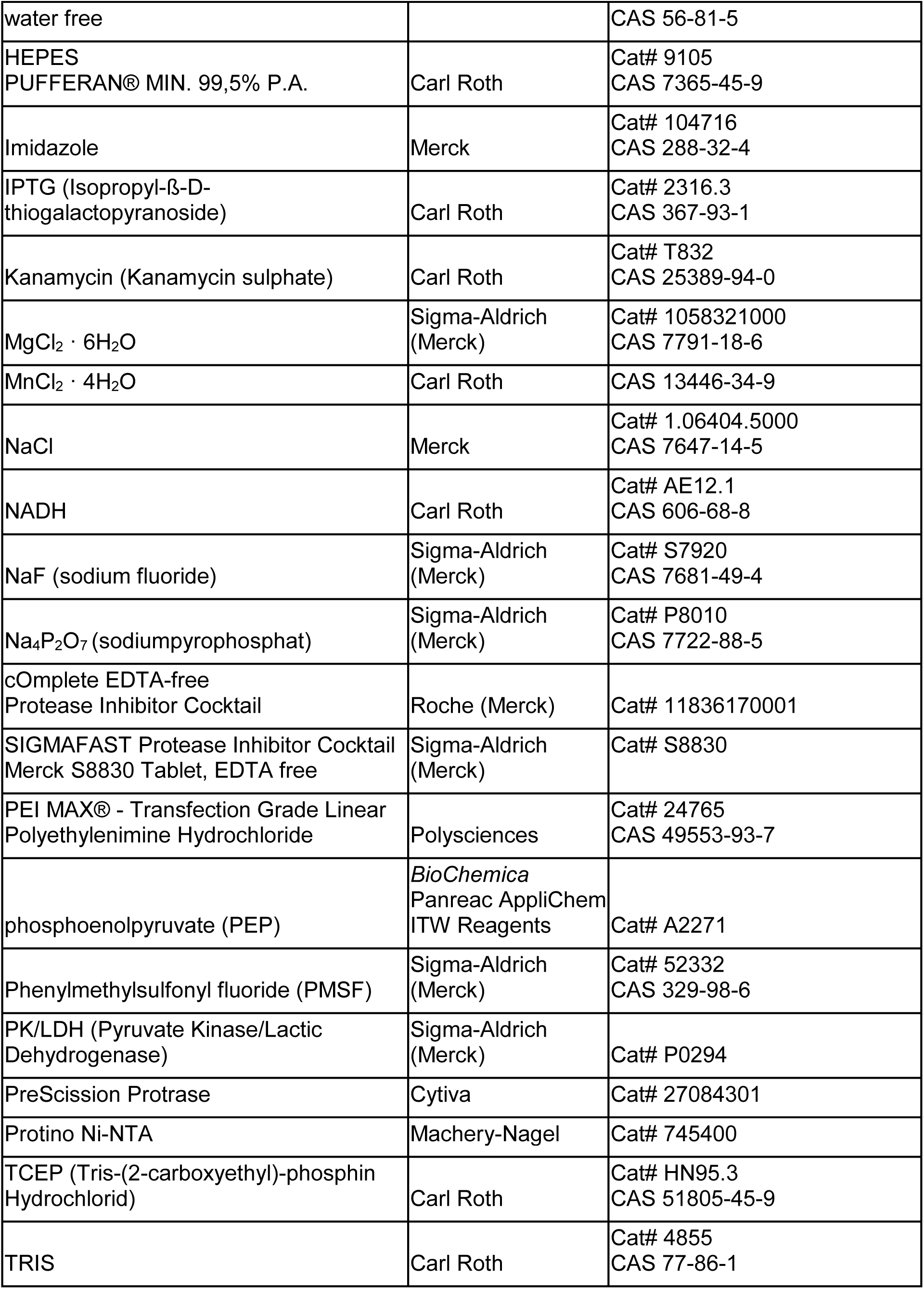
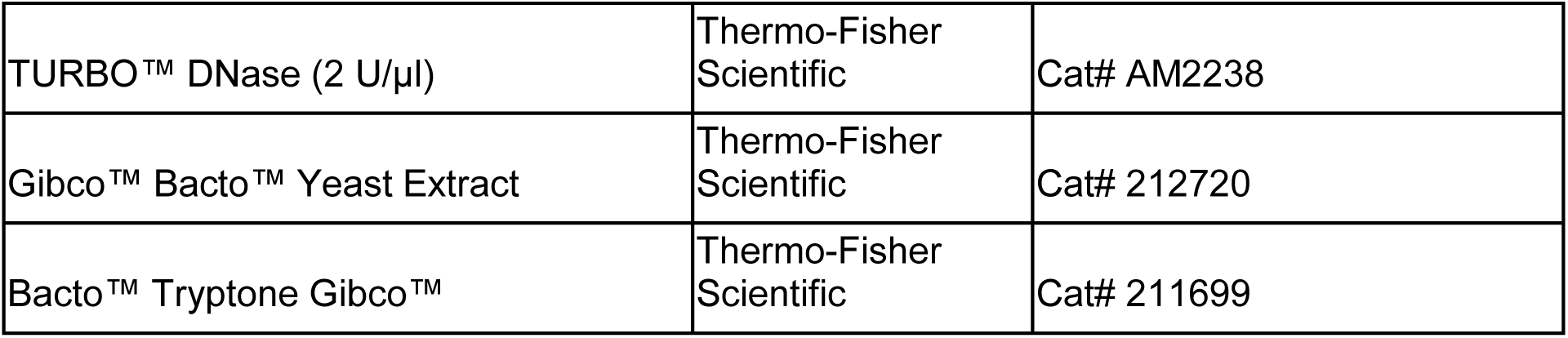

#### Plasmids and recombinant proteins

| Plasmids | Origin | Purpose |
| --- | --- | --- |
| pACEBac1_MRE11_3C_2xFlag_RAD50 | This study | Expression in Hi5 cells |
| pACEBac1_MRE11_3C_2xFlag_RAD50_NBS1 | This study | Expression in Hi5 cells |
| pACEMam1_MRE11<sup>H129N</sup>_3C_2xFlag_RAD50_NBS1 | Ref. <sup>64</sup> | Expression in HEK293T |
| pACEMam1_MRE11_2xFlag_RAD50<sup>apex</sup>_NBS1 | Ref. <sup>64</sup> | Expression in HEK293T |
| pBig1a zz TEV YBBR RAP1 ZZ TEV TRF2 | Ahmet Yildiz, Addgene #185449, RRID:Addgene_185449 | Subcloning into pFastBac-MBP |
| pFastBac-MBP-3C-TRF2-His<sub>10</sub> | This study | Expression in Hi5 cells |
| pET47b_His<sub>6</sub>-SUMO_TRF2<sup>iDDR</sup> (438-485) | This study | Expression in *E.coli* Rosetta |
| pET47b_His<sub>6</sub>-SUMO_TRF2<sup>iDDR-Myb</sup> (438-542) | This study | Expression in *E.coli* Rosetta |

| Recombinant Proteins |  | Purpose |
| --- | --- | --- |
| MRN WT (MRE11-RAD50-NBS1) | This study | EM, biochemistry |
| MR (MRE11-RAD50) | This study | EM, biochemistry |
| M<sup>H129N</sup> RN | Ref. <sup>64</sup> | EM |
| MR<sup>apexN</sup> | Ref. <sup>64</sup> | Crosslinking MS |
| TRF2 full-length | This study | EM |
| TRF2 (aa 438-485) (TRF2<sup>iDDR</sup>) | This study | purification, biochemistry |
| TRF2 (aa 438-542) (TRF2<sup>iDDR-Myb</sup>) | This study | EM, biochemistry |

#### Deposited Maps

| Deposited Maps |  | Identifier |
| --- | --- | --- |
| MR-DNA | This study | EMDB: 52959 (3.21 Å) and PDB: 9Q9H |
| MRN-DNA | This study | EMDB: 52960 (3.08 Å) and PDB: 9Q9I |
| MR-TRF2<sup>iDDR-Myb</sup>-DNA | This study | EMDB: 52962 (2.7 Å) and PDB: 9Q9K |
| MRN-TRF2<sup>iDDR-Myb</sup>-DNA | This study | EMDB: 52961 (3.01 Å) and PDB: 9Q9J |
| MRN-TRF2-DNA | This study | EMDB: 52964 (2.97 Å) and PDB: 9Q9M |

#### Oligonucleotides

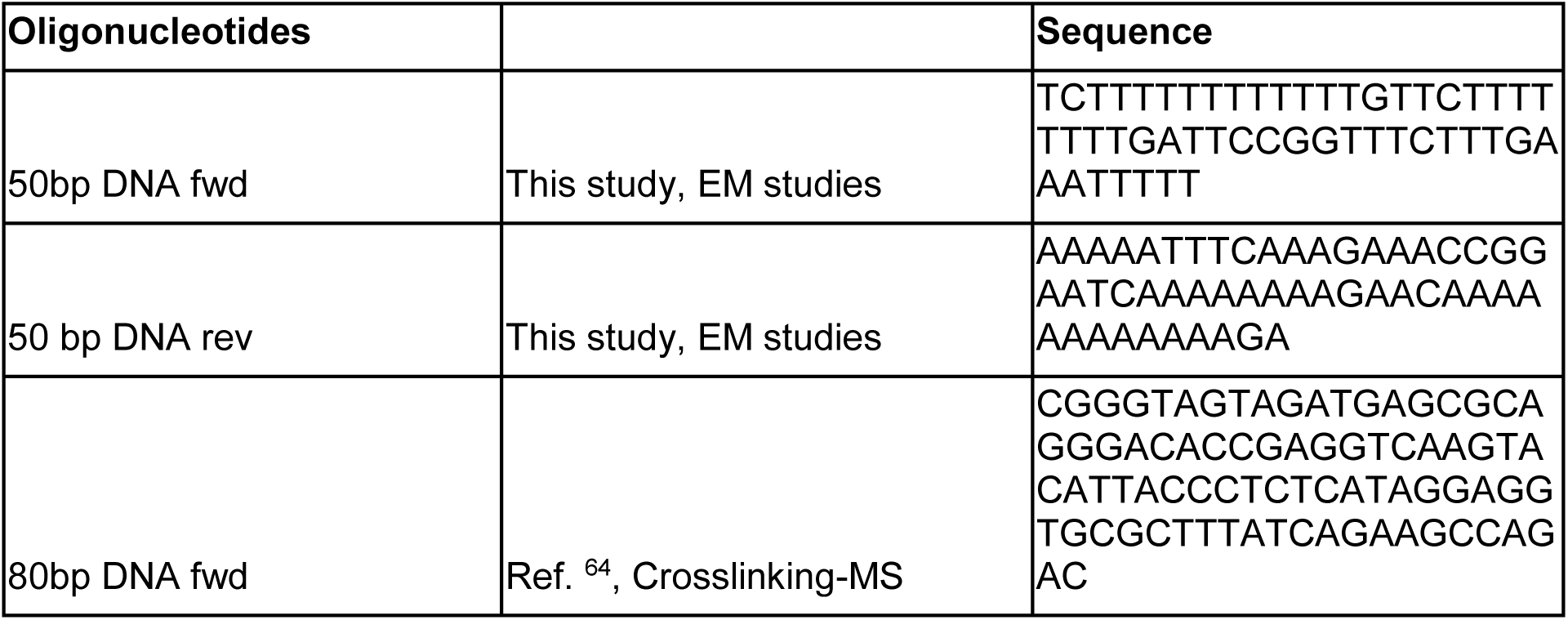
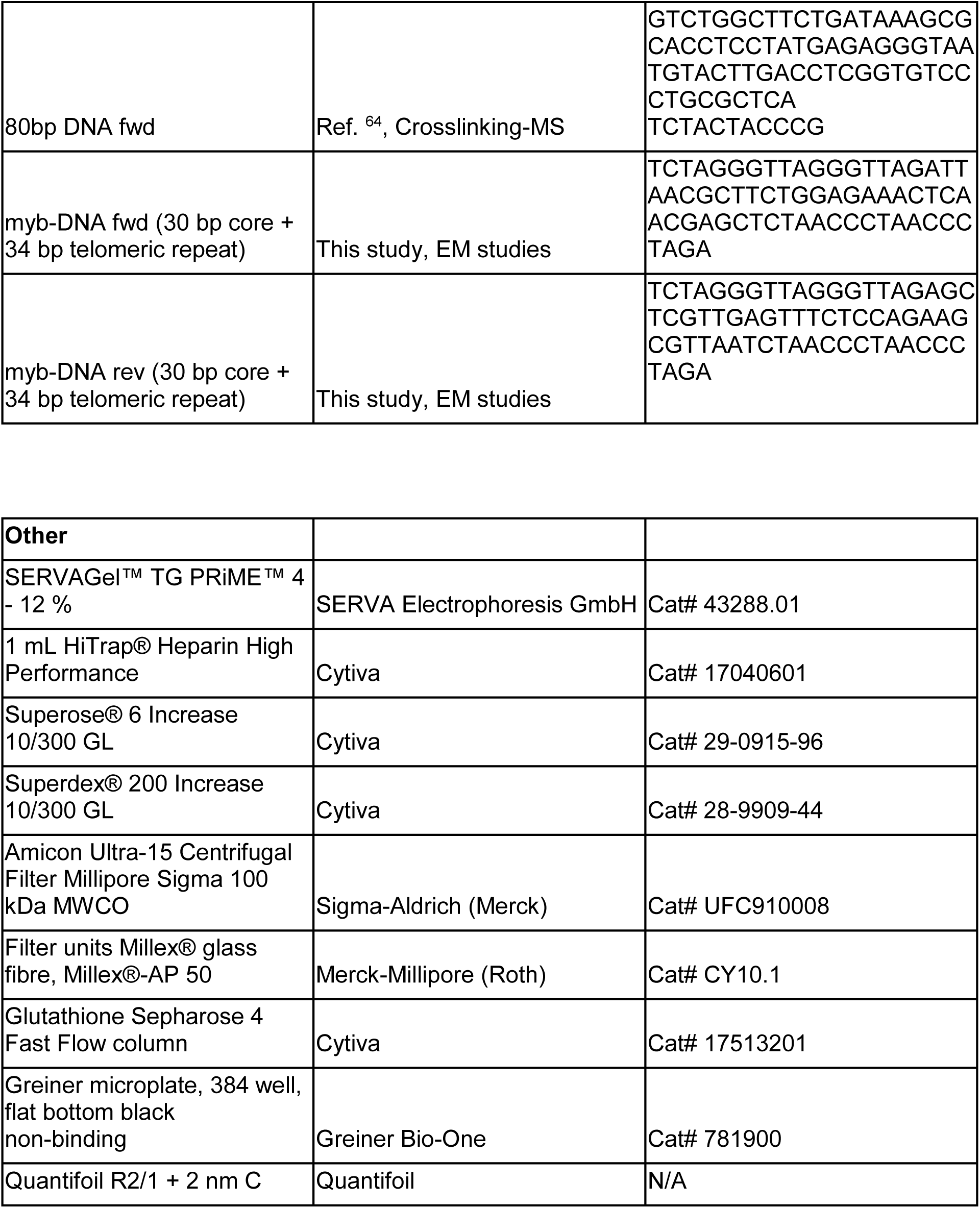

#### Software and algorithms

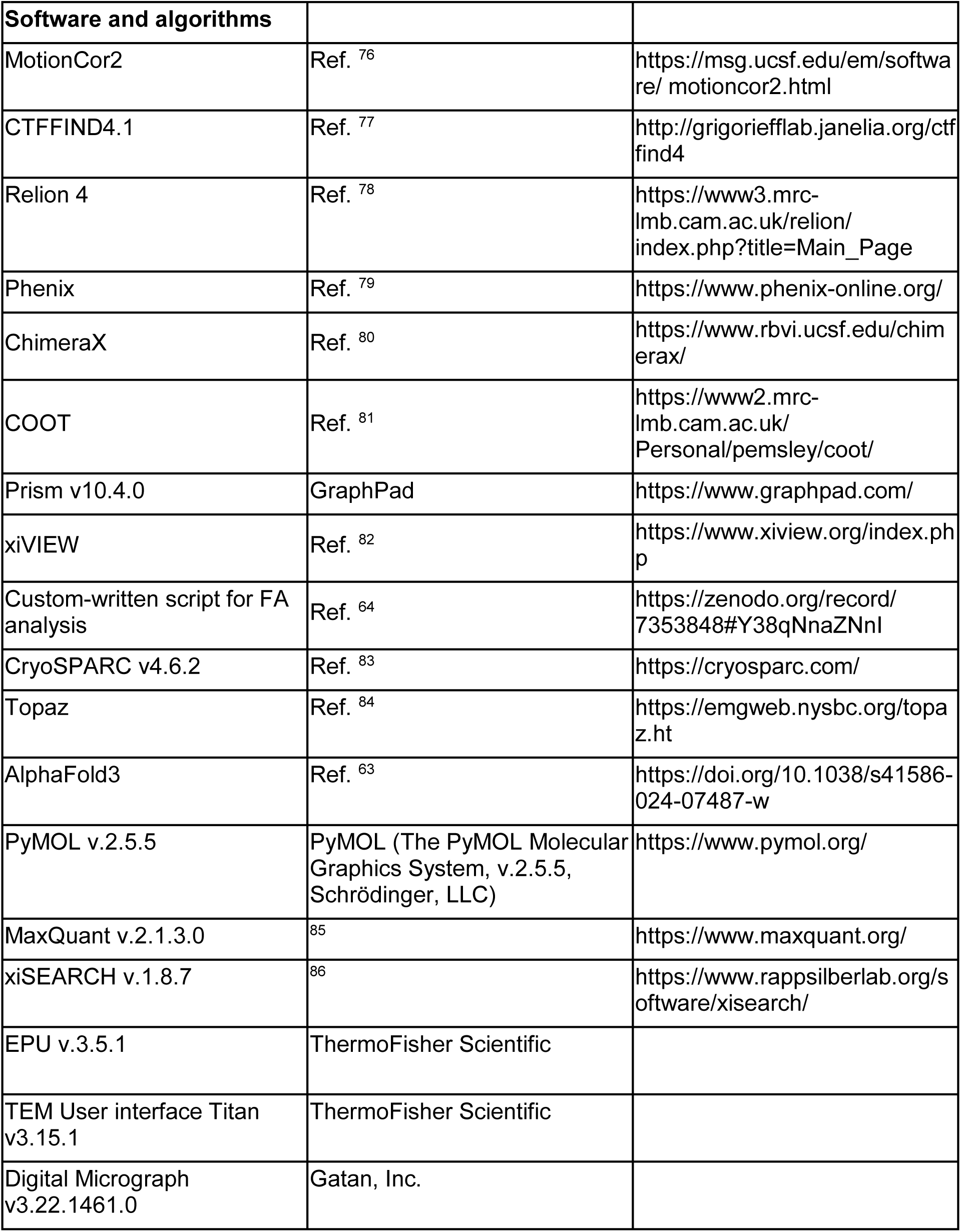

#### Experimental models: Cell lines, Organisms

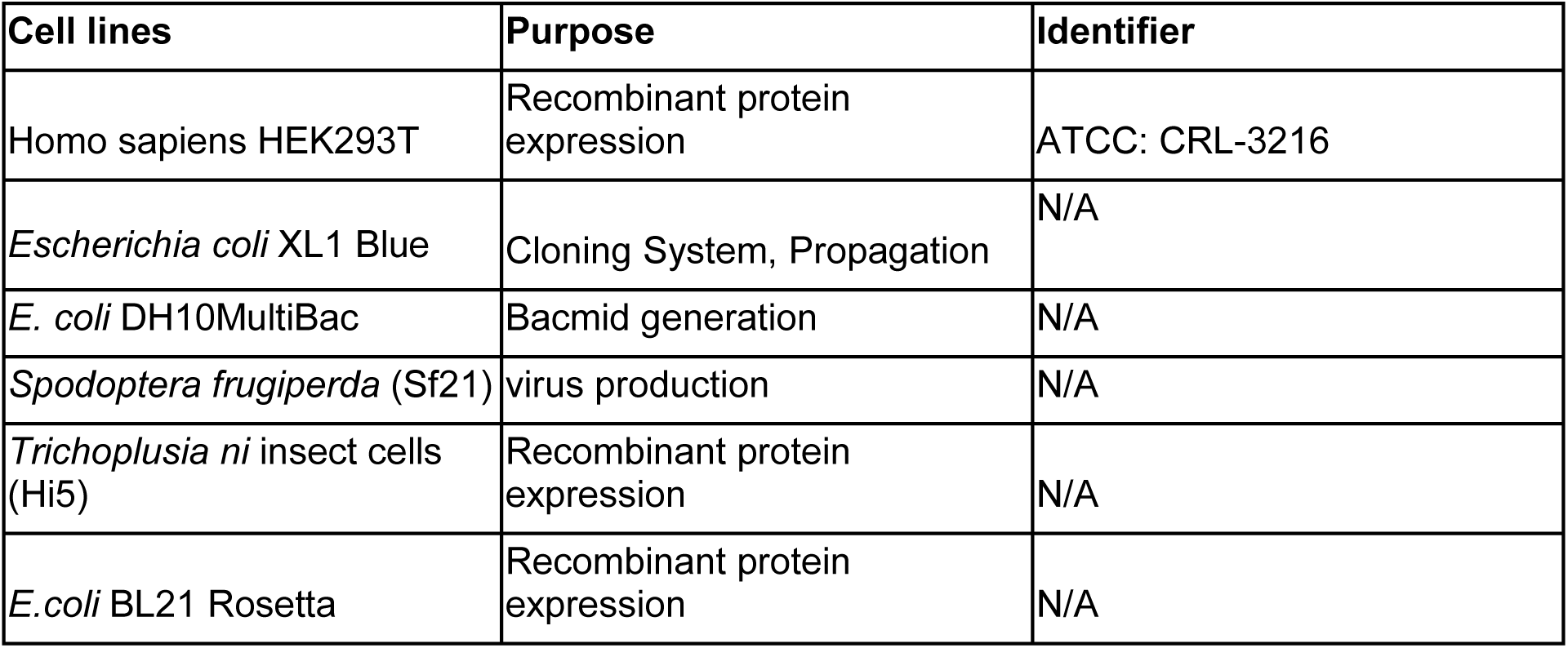

| Cell lines | Purpose | Identifier |
| --- | --- | --- |
| Homo sapiens HEK293T | Recombinant protein expression | ATCC: CRL-3216 |
| *Escherichia coli* XL1 Blue | Cloning System, Propagation | N/A |
| *E. coli* DH10MultiBac | Bacmid generation | N/A |
| *Spodoptera frugiperda* (Sf21) | virus production | N/A |
| *Trichoplusia ni* insect cells (Hi5) | Recombinant protein expression | N/A |
| *E.coli* BL21 Rosetta | Recombinant protein expression | N/A |

#### Organisms as source for materials used in experiments

For the amplification of plasmid DNA, we used *Escherichia coli* XL1 Blue cells and *E. coli* DH10MultiBac cells for bacmid generation. *Spodoptera frugiperda* (Sf21) insect cells were used for virus production. *E.coli* Rosetta2, *Trichoplusia ni* insect cells (Hi5) and *Homo sapiens* HEK293T cells were used for recombinant protein expression.

### Protein purification

#### Expression and purification of MR and MRN complexes

MRN complexes were either expressed in HEK293T cells or *Trichoplusia ni* High Five (Hi5) cells. Purification protocols were adapted from our previously described protocols and applied to purifications from both expression systems^64,87^. In brief: pACEMam1_pMDC_M^H129^RN plasmid was transfected into 500 mL HEK293T cells, cultured for 72 h at 37 °C and harvested by centrifugation (1000 g, 4°C, 10 min.). In case when insect cells were used as the expression system, bacmids were generated in *E.coli* DH10MultiBac as previously described according to the MultiBac system by transformation into *E. coli* DH10MultiBac cells followed by blue-white selection^88^. Baculoviruses containing MRN wild-type or MR were generated in *Spodoptera frugiperda* (SF21) cells. Virus titers were determined by small-scale test expressions. 1 L of *Trichoplusia ni* High Five cells (Invitrogen) seeded to 1 mio/mL were infected with 1:500 of respective baculovirus and cultured for 72 h at 27°C. Cells were harvested by centrifugation (4000 rpm, 4°C, 15 min.). All subsequent purification steps were carried out either in the cold room or on ice. Pellets were homogenized in lysis buffer (50 mM HEPES pH 8.0, 250 mM NaCl, 10% glycerol, 0.1 mM DTT, 0.5 mM MnCl_2_, 1 mM MgCl_2_, two cOmplete EDTA-free Protease Inhibitor Cocktail tablets (Roche), 2 µL TURBO DNase I (Thermo Fisher Scientific), 20 mM NaF and 12 mM Na_2_P_2_O_7_. Lysates were sonicated for 2x 45 sec (40% duty cycle, 5 output control) and centrifuged at 16.000 rom for 1 h at 4°C. Supernatants were filtered through a Millex® glass fibre filter unit, Millex®-AP 50 (Roth). Clarified lysates were applied onto 2 mL FLAG M2 affinity resin (Sigma) in a gravity flow column, equilibrated in low salt buffer (50 mM HEPES pH 8.0, 250 mM NaCl, 10% glycerol, 0.1 mM DTT, 0.5 mM MnCl_2_) and incubated for 2 h rolling. Beads were washed with 10 column volumes (CV) low salt buffer, 5 CV high salt ATP-buffer (50 mM HEPES pH 8.0, 1 M NaCl, 2 mM ATP, 5 mM MgCl_2_ 10% glycerol, 0.1 mM DTT, 0.5 mM MnCl_2_) followed by 5 CV Buffer high salt buffer w/o ATP (50 mM HEPES pH 8.0, 1 M NaCl, 10% glycerol, 0.1 mM DTT, 0.5 mM MnCl_2_). Afterwards, beads were re-equilibrated in low salt buffer. Protein was eluted with 0.2 mg/mL 3x FLAG-peptide (TargetMol) in low salt buffer to a total volume of 5 CV. The eluate was concentrated using a Amicon® Ultra Centrifugal Filter, 100 kDa MWCO and applied to size exclusion chromatography (Superose 6, 10/300, Cytiva) in SEC buffer (25 mM HEPES pH 8.0, 150 mM NaCl, 1 mM DTT). Stoichiometric MRN complex eluted at 0.5 CV. The integrity of the peak fractions was analyzed by SDS-PAGE (SERVA) and confirmed by Mass Photometry (Refeyn), for which proteins were diluted to 50 nM just before the measurement (Fig. S1A,B).

#### Expression and purification of TRF2 complexes

The full-length human TRF2 gene was amplified from Addgene plasmid #185449 shared by Ahmet Yildiz (http://n2t.net/addgene:185449; RRID:Addgene_185449) and subcloned into a pFastBac vector using NheI and XhoI (pFB-MBP-3C-HsTRF2-His_10_). Baculovirus containing TRF2 was generated according to the protocol described for MR/MRN. Expression of full-length TRF2 was carried out in 1 L of Hi5 insect cells. Cells were harvested by centrifugation (500 g, 4°C, 15 min) 72 h post-infection, flash-frozen in liquid nitrogen, and stored at –80 °C.

For purification, 10 mL of cell pellets were thawed and resuspended in 70 mL of lysis buffer (50 mM Tris-HCl, pH 7.5, 1 mM TCEP) supplemented with 1 mM phenylmethylsulfonyl fluoride (PMSF) and one tablet of Sigma protease inhibitor cocktail. The cells were allowed to swell for 20 minutes at 4 °C with gentle stirring. Subsequently, 40 mL of 50 % glycerol and 7.8 mL of 5 M NaCl were added drop-wise while mixing, followed by a 30-minute incubation. The cell suspension was centrifuged at 38.000 g for 45 min at 4°C to obtain clear lysate, which was then filtered through a a Millex® glass fibre filter unit, Millex®-AP 50 (Roth).

The resulting supernatant, supplemented with 10 mM imidazole, was incubated with 1 mL of pre-equilibrated Ni-NTA agarose resin (Machery-Nagel) for 1 h at 4 °C. The resin was washed with 100 CV of high-salt buffer (lysis buffer + 500 mM NaCl + 20–50 mM imidazole) followed by 20 mL of low-salt buffer (lysis buffer + 100 mM NaCl). MBP-TRF2-His_10_ was eluted using 15 CVs of low-salt buffer with 500 mM imidazole. To remove the MBP tag, PreScission protease (1 µg per 50 µg MBP-TRF2-His_10_) was added to the eluate and incubated for 3 h at 4 °C. The TRF2 protein was further purified using a 1 mL HiTrap Heparin column (Cytiva), developed with a NaCl gradient (100–1000 mM) in lysis buffer containing 0.5 mM TCEP and 5 % glycerol. Fractions containing full-length TRF2 were pooled and concentrated to approximately 1 mL before being injected onto a Superdex 200 Increase 10/300 GL column (Cytiva) equilibrated in storage buffer (25 mM HEPES, pH 7.5, 150 mM NaCl, 0.5 mM TCEP, and 5 % glycerol). Purity was assessed via SDS-PAGE. Peak fractions were pooled, concentrated to approximately 12 µM (dimer), aliquoted, and flash-frozen in liquid nitrogen.

TRF2 variants, including TRF2^iDDR^ (438-485) and TRF2^iDDR-Myb^ (438-542), were expressed as His₆-SUMO fusion proteins in *E. coli* Rosetta cells using pET47b vectors. Transformed Rosetta cells were grown in 4 L of 2YT medium (2× Yeast Extract Tryptone) supplemented with kanamycin (50 µg/mL) and chloramphenicol (30 µg/mL) at 37 °C with aeration until the culture reached an OD_600_ of approximately 1.0. Protein expression was induced by adding 0.5 mM IPTG, followed by incubation at 16 °C for 18 h. Cells were harvested (5000 g, 4°C,15 min), flash-frozen in liquid nitrogen, and stored at –20°C.

For purification, the cell pellet was resuspended in 100 mL of lysis buffer supplemented with 500 mM NaCl, 1 mM PMSF, and one Sigma protease inhibitor cocktail tablet. Cells were lysed via sonication, and the lysate was clarified by centrifugation (38.000 g, 45 min, 4°C). The supernatant, supplemented with 10 mM imidazole, was applied via gravity flow to 6 mL of pre-equilibrated Ni-NTA agarose resin (Machery-Nagel). The column was washed with 15 CV of high-salt buffer (lysis buffer + 1 M NaCl + 20 mM imidazole), followed by 8 CV of low-salt buffer (lysis buffer + 100 mM NaCl + 40 mM imidazole). The TRF2 variants were eluted using 3 CV of low-salt buffer supplemented with 500 mM imidazole. To remove the His₆-SUMO fusion tag, GST-tagged PreScission protease was added to the eluate (1:50 enzyme-to-protein ratio), and the mixture was dialyzed overnight at 4°C against 1 L of buffer (50 mM Tris-HCl, pH 7.5, 100 mM NaCl, 1 mM TCEP, and 1 mM EDTA). The His₆-SUMO tag and uncleaved fusion protein were removed via a 6 mL Ni-NTA column, while PreScission protease was eliminated using a 2 mL Glutathione Sepharose 4 Fast Flow column. Final purification was performed using a Superdex 200 Increase 10/300 GL column (Cytiva) equilibrated in storage buffer (25 mM HEPES, pH 7.5, 150 mM NaCl, 1 mM TCEP, and 1 mM EDTA). Protein purity was confirmed via SDS-PAGE. Peak fractions were pooled, concentrated to approximately 417 µM, aliquoted, and flash-frozen in liquid nitrogen.

#### DNA substrates

The oligonucleotides used in this study were acquired from Metabion (Germany). Modified sequences (6-FAM labeled) were obtained HPLC-purified and lyophilized. Complementary ssDNA sequences were annealed to one another in a PCR cycler starting at 95°C with gradually decreasing temperatures (0.1°C/sec.) down to 25°C. For annealing, an annealing buffer was used (25 mM TRIS pH 7.5, 50 mM NaCl, 10 mM MgCl_2_). In case of modified sequences, the complementary sequence was used in 1.1x excess compared to the labeled/modified oligonucleotide. dsDNA was either immediately used or stored at -20°C.

#### Fluorescence anisotropy DNA binding assays

Fluorescence polarization anisotropy (FA) was used to monitor MR/MRN and TRF2 binding to 30bp-Myb dsDNA. 30 nM of 6-FAM-labelled DNA was incubated with increasing amounts of MR(N) and/or TRF2/MRN in assay buffer (25 nM HEPES 8.0, 100 mM NaCI, 5 mM MgCI2, 0.2 mg/ml BSA, 1 mM DTT). 1 mM ATP was added if not stated otherwise. When both MRN and TRF2 present in the reaction, a constant concentration of TRF2 (1.6 μM) was used with increasing amounts of MRN. FA was measured at an excitation wavelength of 488 nm and an emission wavelength of 520 nm using an automated polarization microscope^89,90^. Measurements were taken every 5 min for 30 min to monitor MRN-DNA binding stability. In each well, twelve different z-planes were measured to reduce the possibility of erroneous FA values caused by potential fluorescing protein-DNA aggregates. The reactions were prepared and measured in triplicates. The median FA values were calculated as described in ref.^64^. The final FA value of each sample was the average of median FA values over the time course of 30 min.

#### NADH-coupled ATPase assay

ATPase rates of MRN, MR and complex mutants were assessed using a plate-reader based ATPase assay, that is coupled to NADH oxidation. All steps prior to the measurement were carried out on ice. 300 nM protein (MR and MRN WT from Hi5 cells) was incubated with assay buffer (100 mM NaCl, 25 mM HEPES pH 7.5, 2 mM MgCl_2_, 1 mM MnCl_2_, 1 mM phosphoenolpyruvate (PEP), 0.15 mM ATP, 25 U/mL Lactate Dehydrogenase/Pyruvate kinase (Sigma-Aldrich), 0.1 mM NADH, 0.1 mg/mL BSA (NEB), 0.5 mM DTT) in 50 µL final reaction volume. To stimulate ATPase activity, substrate (50 bp DNA in case of MR and MRN measurements; 30 bp Myb-DNA in case of TRF2^iDDR-Myb^ measurements) was added to achieve 1000 nM final DNA concentration. Decreasing NADH concentrations correlate with fluorescence decay of NADH which was monitored over 60 (MR/MRN + DNA) or 90 min (MR/MRN + TRF2) at 37°C in a Greiner microplate, 384 well, flat bottom, non-binding black 384 plate (Greiner Bio-One) using 340 nm laser for excitation and emission at 460 nm in a Tecan Spark plate reader (Tecan). The reactions were prepared and measured in triplicates unless otherwise noted (only duplicate in case of MR+TRF2). The ATP turnover was determined using steady-state rate at maximal initial linear rates and corrected subtracting buffer blank.

#### Cryo-EM grid preparation

Freshly purified MR/MRN was directly used for cryo-EM sample preparation. The proteins were diluted to final concentrations of 0.4 µM (∼0.2 mg/mL) in 25 mM HEPES pH 7.5, 120 mM NaCl, 5 mM MgCl_2_, 1 mM MnCl_2_, 1 mM DTT, 1 mM ATP and incubated for 10 min. at room temperature. DNA substrate (50 bp see Table Oligonucleotides) was added to reach a final concentration of 0.3 µM and incubated at 35°C for at least 30 min followed by the addition of 1 mM BeF_x_. In case TRF2 full-length or TRF2^iDDR^ was added, DNA containing telomeric repeat regions (referred to as 30 bp Myb-DNA, see Table Oligonucleotides) and TRF2 were pre-incubated for 10 min at 35°C, then added to the MR(N) complex. Grids were prepared using a Leica EM GP plunge freezer (Leica) at 12°C and 95% humidity. Just before plunging, the reaction was supplemented with octyl β-D-glucopyranoside (β-OG) at final concentration of 0.05%. 4.5 µL of sample were applied onto a glow-discharged Quantifoil® R2/1 + 2nm carbon Cu 200 grid. The samples were pre-blotted for 20 sec, blotted for 2.3 sec, before vitrification in liquid ethane.

#### Cryo-EM data acquisition collection

MR-DNA dataset was collected on FEI Titan Krios G3 transmission electron microscope (300 kV) equipped with a GIF quantum energy filter (slit width 20 eV) and a Gatan K2 Summit direct electron detector (software used: EPU 3.5.1, TEM User interface Titan 3.15.1, Digital Micrograph 3.22.1461.0). All other datasets were collected on FEI Titan Krios G3 transmission electron microscope (300 kV) equipped with a Selectris X imaging filter (slit width 10 eV) and a Falcon 4 direct electron detector (software used: EPU 3.5.1, TEM User interface Titan 3.15.1).

For the MR-DNA dataset, 4047 movies were collected at a nominal magnification of 130,000x (1.06 Å/pix), a defocus range of -1.1 to -2.9 µm, and a total electron dose of 42.377 e−/Å2 fractionated into 40 frames over 10 sec. The MRN-DNA dataset (16,890 movies), MR-TRF2^iDDR-Myb^-DNA dataset (18,134 movies), MRN-TRF2^iDDR-Myb^-DNA dataset (33,072 movies), MRN-TRF2-DNA dataset (39,320 movies) were collected at a nominal magnification of 165,000x (0.727 Å/pix), a defocus range of -0.5 to -2.6 µm, and a total electron dose of 40 e−/Å2 fractionated into 40 movie frames over 2.75 sec.

#### Cryo-EM image processing

Movie frames were motion corrected using MotionCor2 1.4.5^76^. All subsequent cryo-EM data processing steps were carried out using CryoSPARC (v4.6.2 and former versions). The CTF parameters of the datasets were determined using patch CTF estimation job integrated in the CryoSPARC software framework^83^. The masks for the calculation of FSC were created by CryoSPARC automasking. The resolutions reported are calculated based on the gold-standard Fourier shell correlation criterion (FSC = 0.143) with 3D FSC plots using Remote 3DFSC Processing Server^91^. The exact processing schemes are depicted in Supplementary Figs 2,3,7,8 and 9. Data collection and refinement statistics are summarized in Supplementary Table 1.

For the MR-DNA structure, particles were initially picked on 4,047 micrographs using a blob picker with an applied diameter of 120 to 360 Å (Supplementary Fig.2). Picked particles were extracted with a box size of 360 pixel, and subjected to multiple rounds of 2D classification, resulting in an initial cleaned subset of 2D classes comprising 573,534 particles. The 2D classes with clearly defined features were selected and served as templates for a reference-based template picking approach, and subsequently as input for Topaz train^84^. A total pool of 817,041 particles was further sorted by ab initio reconstruction and heterogeneous refinement. The yielded class of 301,918 particles with clearly defined features were further sorted by rounds of 2D classification, homogeneous refinement and non-uniform refinement, leaving a new particle set of 226,766 particles. 3D classification was applied to separate conformational states of MR with symmetrically attached MRE11 and partially detached MRE11. The final resolution of MR-DNA reconstruction (95,397 particles), after homogeneous refinement and non-uniform refinement of the former class, was 3.21 Å. The sharpened map of this reconstruction was used further for model building and refinement.

For the MRN-DNA structure, particles were initially picked on 16,890 micrographs using a blob picker with an applied diameter of 120 to 360 Å (Supplementary Fig.3). Picked particles were extracted with a box size of 360 pixel, and subjected to multiple rounds of 2D classification, resulting in an initial cleaned subset of 2D classes comprising 573,534 particles. Reasonable 2D classes were selected and served as templates for a reference-based template picking approach, and subsequently as input for Topaz train^84^. A total pool of 929,509 particles was further sorted by 2D classification to remove a subset of 206,194 poorly resolved particles. The remaining 723,315 particles were subjected to ab-initio reconstruction and heterogeneous refinement. Particles in reasonable 3D volumes were further sorted by homogeneous refinement and 3D classification, which allowed separation of two well-resolved states of MRN-DNA: MRN with symmetrically attached MRE11 (175,312 particles) and partially detached MRE11 (204,607 particles). 3D classification and homogeneous refinement were further applied to the former class, leading to the final MRN-DNA reconstruction at 3.08 Å using 172,784 particles. The sharpened map of this reconstruction was used further for model building and refinement.

For the MR-TRF2^iDDR-Myb^-DNA structure, particles were initially picked on 18,134 micrographs using a blob picker with an applied diameter of 120 to 360 Å (Supplementary Fig.7). Picked particles were extracted with a box size of 360 pixel, and subjected to multiple rounds of 2D classification, resulting in an initial cleaned subset of 2D classes comprising 599,266 particles. Reasonable 2D classes were selected and served as templates for a reference-based template picking, and subsequently as input for Topaz train^84^. A total pool of 1,109,607 particles were subjected homogeneous refinement and 3D classification, which allowed separation of two well-resolved states of MRN-DNA: MRN with symmetrically attached MRE11 (369,251 particles) and partially detached MRE11 (319,438 particles). Particles from the former class were further sorted by homogeneous refinement, non-uniform refinement and 3D classification. The final resolution of MR-TRF2^iDDR-Myb^-DNA reconstruction after non-uniform refinement was 2.7 Å containing 274,928 particles. The sharpened map of this reconstruction was used further for model building and refinement. Particles from the MRE11-partially-detached class were further processed by 2D and 3D classification, followed by iterative rounds of homogeneous refinement. The resulting subset of 203,202 particles was used to reconstruct the cryo-EM map of MR-TRF2^iDDR-Myb^ -DNA (MRE11-detached). The sharpened map of this reconstruction was used further for model building and refinement. This map and model were only used for the preparation of the figures.

For the MRN-^TRF2iDDR-Myb^-DNA structure, particles were initially picked on 33,072 micrographs using a blob picker with an applied diameter of 120 to 360 Å (Supplementary Fig.8). Picked particles were extracted with a box size of 360 pixel, and subjected to multiple rounds of 2D classification, resulting in an initial cleaned subset of 2D classes comprising 429,260 particles. Reasonable 2D classes were selected and served as templates for a reference-based template picking, and subsequently as input for Topaz train^84^. A total pool of 725,737 particles were subjected homogeneous refinement and 3D classification, which allowed separation of two well-resolved states of MRN-TRF2iDDR-Myb-DNA: MRN with symmetrically attached MRE11 (292,228 particles) and partially detached MRE11 (146,293 particles). Particles from the former class were further sorted by homogeneous refinement and 3D classifications, followed by non-uniform refinement. The final resolution of MRN-TRF2^iDDR-Myb^-DNA reconstruction after homogeneous refinement was 3.01 Å (129,300 particles). The sharpened map of this reconstruction was used further for model building and refinement.

For the MRN-TRF2-DNA structure, particles were initially picked on 39,320 micrographs using a blob picker with an applied diameter of 120 to 360 Å (Supplementary Fig.9). Picked particles were extracted with a box size of 360 pixel, and subjected to multiple rounds of 2D classification, resulting in an initial cleaned subset of 2D classes comprising 435,078 particles.

Reasonable 2D classes were selected and served as templates for a reference-based template picking, and subsequently as input for Topaz train^84^. A total pool of 1,155,493 particles were subjected homogeneous refinement and 3D classification, which allowed separation of two well-resolved states of MRN-TRF2-DNA: MRN with symmetrically attached MRE11 (553,435 particles) and partially detached MRE11 (159,492 particles). Particles from the former class were further sorted by rounds of homogeneous refinement and 3D classification. The resulting subset of 208,916 particles was subjected to a last round of non-uniform refinement, and used to reconstruct the cryo-EM map of MRN-TRF2-DNA at 3.01 Å. The sharpened map of this reconstruction was used further for model building and refinement.

#### Model building

All EM structures discussed in this study were constructed based on an atomic model of the MRN-DNA complex predicted by AlphaFold3. This model was initially docked as a rigid body into the refined 3D reconstruction maps using ChimeraX, followed by manual adjustments and real-space refinement in COOT to achieve the best fit to the density. The bound TRF2 fragment was built *de novo*, guided by prominent side chains and secondary structure predictions from AlphaFold3. Due to map quality limitations, the double-stranded DNA sequences in all structures were modeled as AT-pair oligos. The presence of BeF_x_ and ADP in the nucleotide-binding pocket of RAD50 was confirmed by well-defined densities in the highest-resolution EM map. The models underwent iterative cycles of real-space refinement and manual adjustments using PHENIX and COOT, resulting in excellent stereochemistry as validated by MolProbity. All figures were generated using PyMOL (The PyMOL Molecular Graphics System, v.2.5.5, Schrödinger, LLC) and ChimeraX.

#### Crosslinking mass spectrometry

MRN (produced in HEK293T cells, variant MR^apex^N) was used for crosslinking using BS3 (bis(sulfosuccinimidyl)suberate)) crosslinker (Thermo Fisher Scientific). MRN was incubated with reaction buffer to achieve final concentrations of 300 nM protein and 500 mM 80 bp dsDNA in 25 mM HEPES pH 7.5, 107 mM NaCl, 5 mM MgCl_2_, 1 mM MnCl_2_, 1 mM ATP, 1 mM DT) and ATM (purified according to ^87^) was added to the MRN complex for 30 min at 30°C. BS3 crosslinker was reconstituted freshly according to the manufacturers protocol (147 µM final BS3 in the sample) and incubated for 20 min. The reaction was quenched with 1mM Tris-HCl pH 7.5, supplemented with 4x SDS dye and analyzed on a SERVAGel™ TG PRiME™ 4 - 12 %. The gel was stained with Coomassie dye.

#### In-gel digestion

The excised Coomassie-stained gel was cut into small cubes, followed by destaining in 50% ethanol/25 mM ammonium bicarbonate. Proteins were then reduced in 10 mM DTT at 56°C and alkylated by 50 mM iodoacetamide in the dark at room temperature. Afterwards, proteins were digested by trypsin (1 µg per sample) in 50 mM ammonium bicarbonate at 37°C overnight. Following peptide extraction sequentially in 30% and 100% acetonitrile, the sample volume was reduced in a centrifugal evaporator to remove residual acetonitrile. The peptides were then acidified with formic acid and purified via solid phase extraction in C_18_ StageTips ^92^.

#### Liquid chromatography tandem mass spectrometry

Peptides were analyzed on an Orbitrap Exploris 480 mass spectrometer (Thermo Fisher Scientific) coupled to an EASY-nLC 1200 UHPLC system (Thermo Fisher Scientific). Peptides were separated in an in-house packed 55-cm analytical column (inner diameter: 75 μm; ReproSil-Pur 120 C18-AQ 1.9-μm silica particles, Dr. Maisch GmbH) by online reversed phase chromatography through a 90-min gradient of 2.4-32% acetonitrile with 0.1% formic acid at a nanoflow rate of 250 nl/min. The eluted peptides were sprayed directly by electrospray ionization into the mass spectrometer. Each sample was injected twice and measured using two different combinations of collision energies in stepped mode^93^. Mass spectrometry was conducted in data-dependent acquisition mode using a top10 method with one full scan (resolution: 60,000, scan range: 300-1650 m/z, target value: 3 × 10^6^, maximum injection time: 60 ms) followed by 10 fragment scans via higher energy collision dissociation (HCD; normalized collision energy in stepped mode: 25, 30, 35% or 27, 30, 33%; resolution: 30,000, target value: 1 × 10^5^, maximum injection time: 60 ms, isolation window: 1.4 m/z). Only precursor ions of +3 to +8 charge state were selected for fragment scans. Additionally, precursor ions already isolated for fragmentation were dynamically excluded for 25 s.

#### Mass spectrometry data analysis

Raw data files were pre-processed using the MaxQuant software (version 2.1.3.0)^85^ as previously described^94^ ignoring the de-noising filter. The peak lists (*.HCD.FTMS.sil0.apl files) were searched using xiSEARCH (version 1.8.7)^86^ against a target-decoy database consisting of the protein sequences of the MRN complex and the ATM. Trypsin/P specificity was assigned. Up to 4 missed cleavages were allowed. Crosslink search was based on the BS3 specificity linking K, S, T, Y residues and the protein N-terminus. Carbamidomethyl on cysteine was assigned as fixed modification. Variable modifications included methionine oxidation, BS3 mono-link with a hydrolyzed or Tris-quenched end and loop link. Mass tolerance was 5 ppm at the MS1 level and 6 ppm at the MS2 level. Residue pairs-level FDR was set to 1%.

#### Visualization of Crosslinks

The processed data and the coordinates of the MRN-DNA structure were imported to XiVIEW Ref.^82^ to visualize crosslinks between the MRE11, RAD50, NBS1 subunits by mapping the links onto the EM-structure. The False Discovery Rate (FDR) was set to 1% and the crosslinking distance cutoff to < 30 Å. The match score was adjusted to minimum 10 and maximum 20. The circular crosslinking map was generated in XiVIEW along with the structural map comprising the crosslinks, which was exported to ChimeraX.

### QUANTIFICATION AND STATISTICAL ANALYSIS

To analyze ATPase rates, the linear fluorescence decrease between 25 and 90 min was fitted to a linear regression in Prism (GraphPad) and the slope was used to determine the ATPase rate. For the exact calculation of Data in Fig 1B, Fig 4E, we used the following formula:

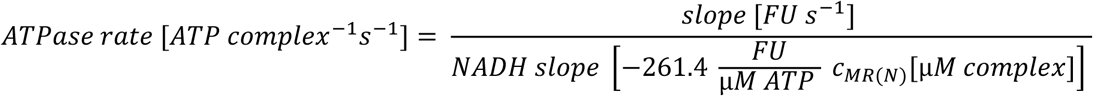

The NADH slope was calculated from a calibration curve that was recorded by carrying out a titration of varying ADP concentrations ranging from 0 to 100 µM in 10 µM increments to constant NADH concentration in the solution lacking protein and DNA.

FA data of MR/MRN binding to 30bp-Myb DNA was analyzed using Prism (GraphPad) by fitting the anisotropy data to a One site Specific binding with Hill slope model, where the following formula was employed (Fig. 2C, Supplementary Fig. 10B):

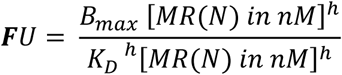

FA data of TRF2 binding to 30bp-Myb DNA was analyzed using Prism (GraphPad) by fitting the anisotropy data to a Two sites Specific binding model, where the following formula was employed (Supplementary Fig. 10A):

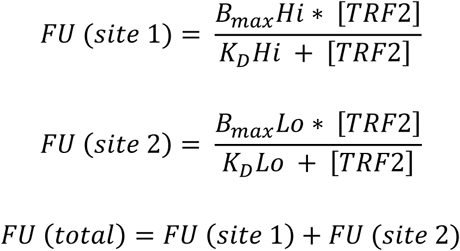

## Supplementary Figure Legends

**Supplementary Figure 1:**
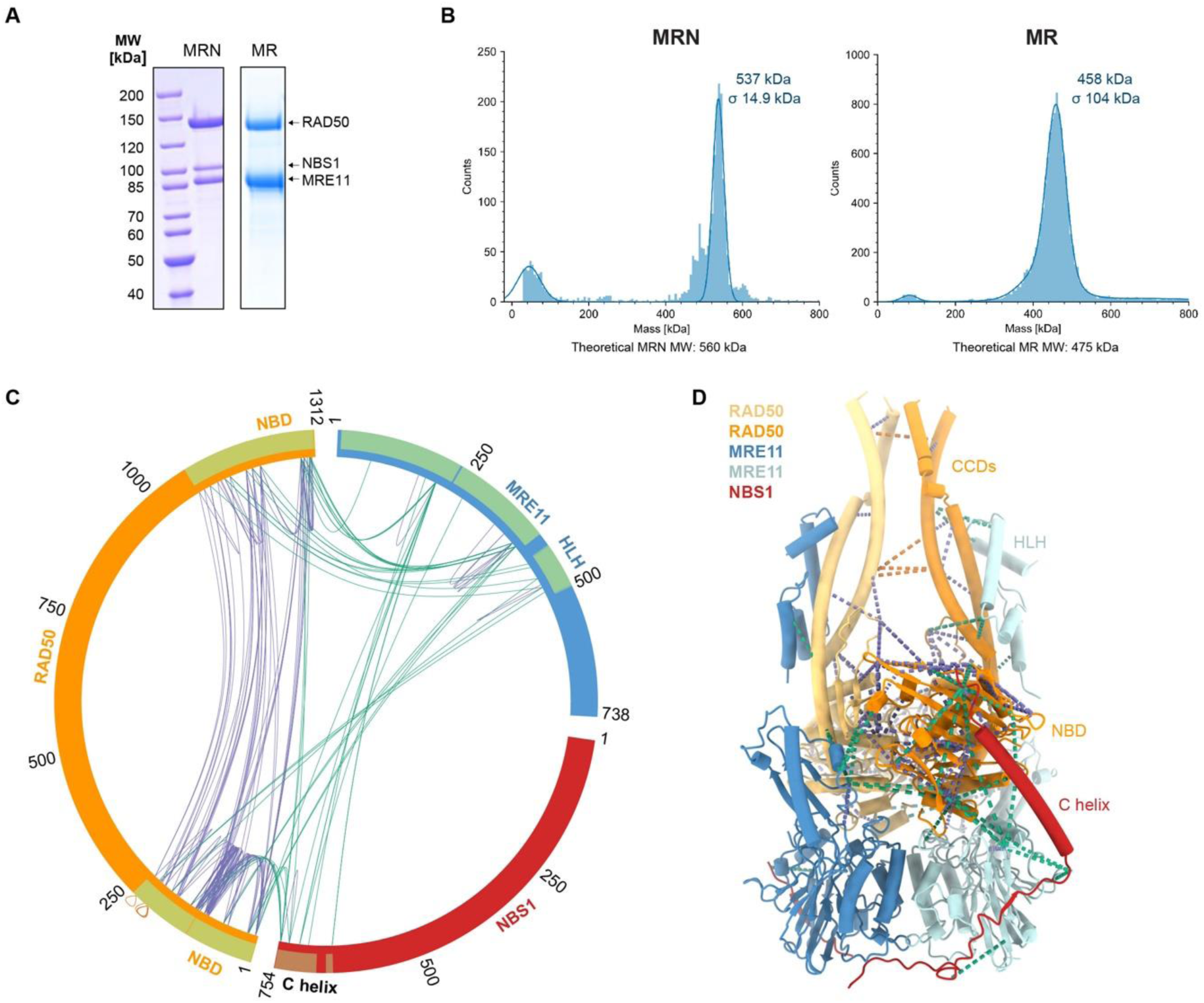
MR(N) complex and chemical cross-linking coupled with mass spectrometry. **A,** Representative SDS-PAGE analysis of purified MR/MRN complexes after gel filtration. **B,** Mass photometry histograms of the MR/MRN complexes (50 nM). **C,** XiVIEW-generated circular crosslinking map of the MRE11, RAD50, NBS1 subunits. The False Discovery Rate (FDR) was set to 1% and the crosslinking distance <30 Å. The PDB annotated regions in each subunit are highlighted in light green. Purple and green lines denote self and heteromeric crosslinks, respectively. **D,** Crosslinks mapped onto the MRN-DNA structure. DNA is not shown for better visual representation.

**Supplementary Figure S2:**
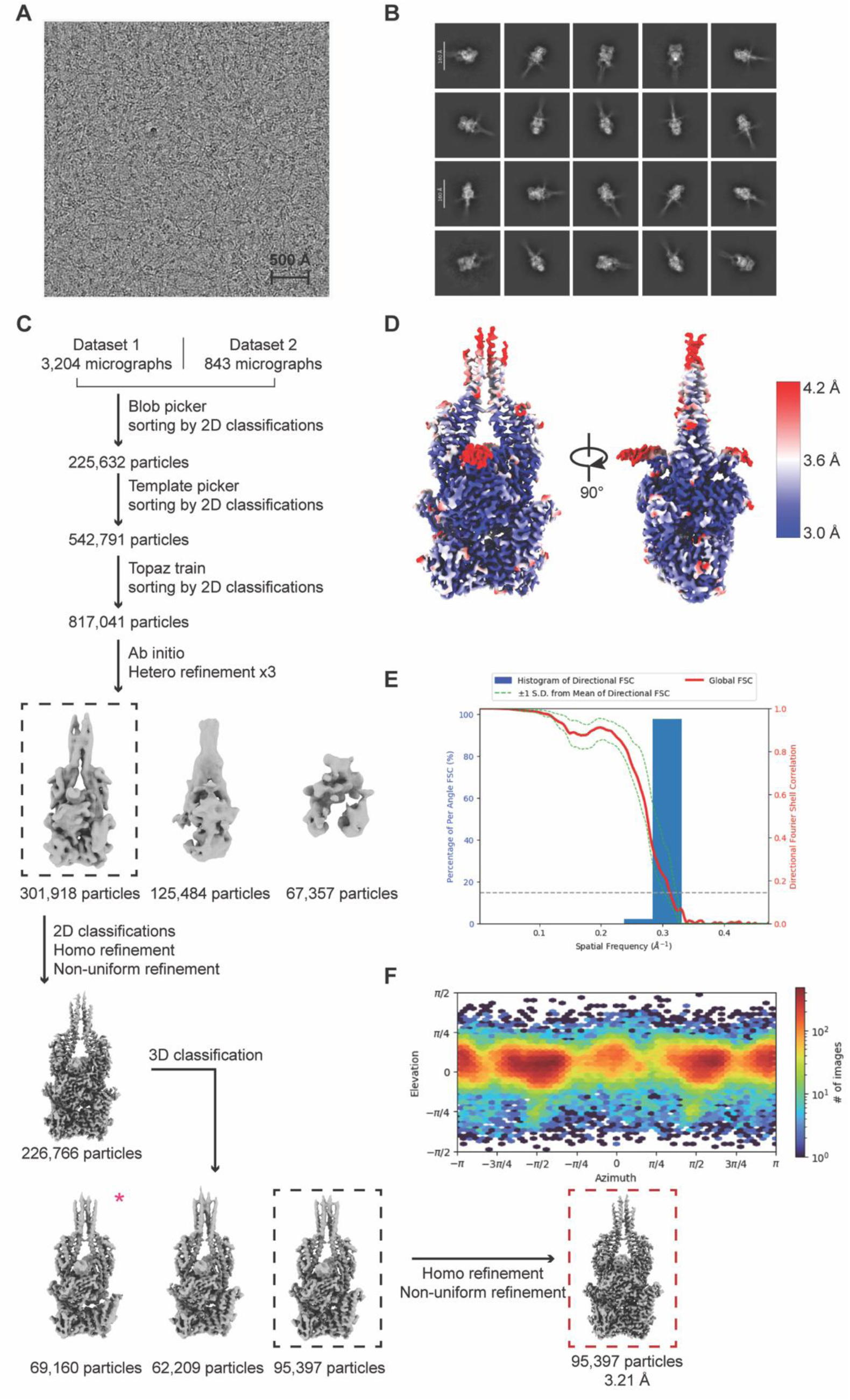
Cryo-EM data analysis of MR-DNA. **A,** Representative micrograph of MR-DNA among 4047 collected movies. **B,** Representative 2D classes of the particles used for the final MR-DNA reconstruction. **C,** Cryo-EM data processing workflow of MR-DNA using cryoSPARC^83^. Pink asterisk denotes the class with MRE11 detached from RAD50^NBD^**. D,** Local resolution visualisation of MR-DNA calculated in cryoSPARC. Blue indicates higher resolution, red indicates lower resolution. **E,** Histogram of directional Fourier shell correlation (FSC) (blue) and global FSC curve (red) of the final MR-DNA reconstruction. The spread of directional resolution values (green dashed lines) is defined as ± 1σ. The grey dashed line shows the 0.143 cut-off criterion, indicating a nominal resolution of 3.21Å. **F,** Angular distribution of the particles used for final MR-DNA reconstruction.

**Supplementary Figure S3:**
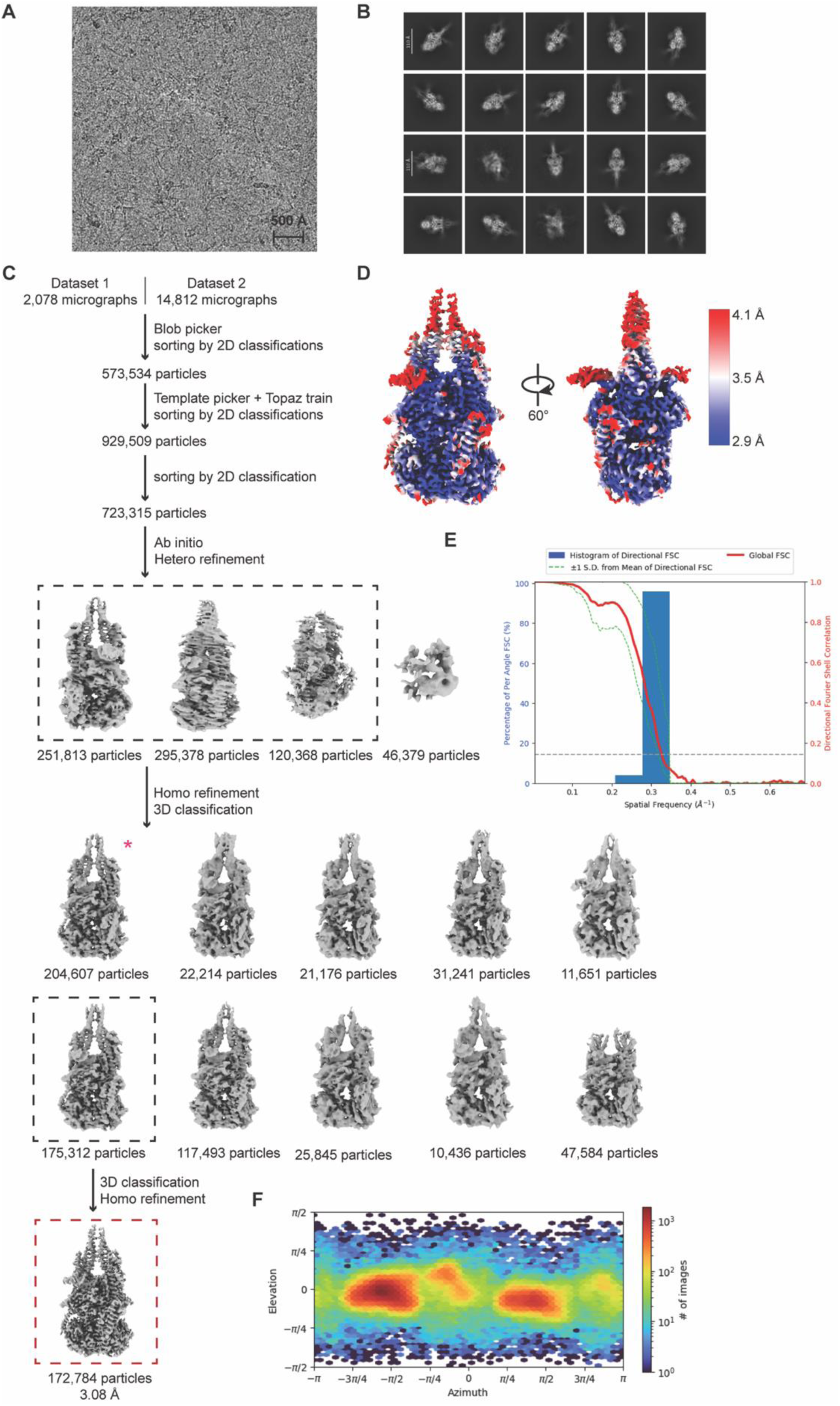
Cryo-EM data analysis of MRN-DNA. **A,** Representative micrograph of MRN-DNA among 16,890 collected movies. **B,** Representative 2D classes of the particles used for the final MRN-DNA reconstruction. **C,** Cryo-EM data processing workflow of MRN-DNA using cryoSPARC^83^. Pink asterisk denotes the class with MRE11 detached from RAD50^NBD^. **D,** Local resolution visualisation of MRN-DNA calculated in cryoSPARC. Blue indicates higher resolution, red indicates lower resolution. **E,** Histogram of directional FSC^91^ (blue) and global FSC curve (red) of the final MRN-DNA reconstruction. The spread of directional resolution values (green dashed lines) is defined as ± 1σ. The grey dashed line shows the 0.143 cut-off criterion, indicating a nominal resolution of 3.08Å. **F,** Angular distribution of the particles used for final MRN-DNA reconstruction.

**Supplementary Figure S4.**
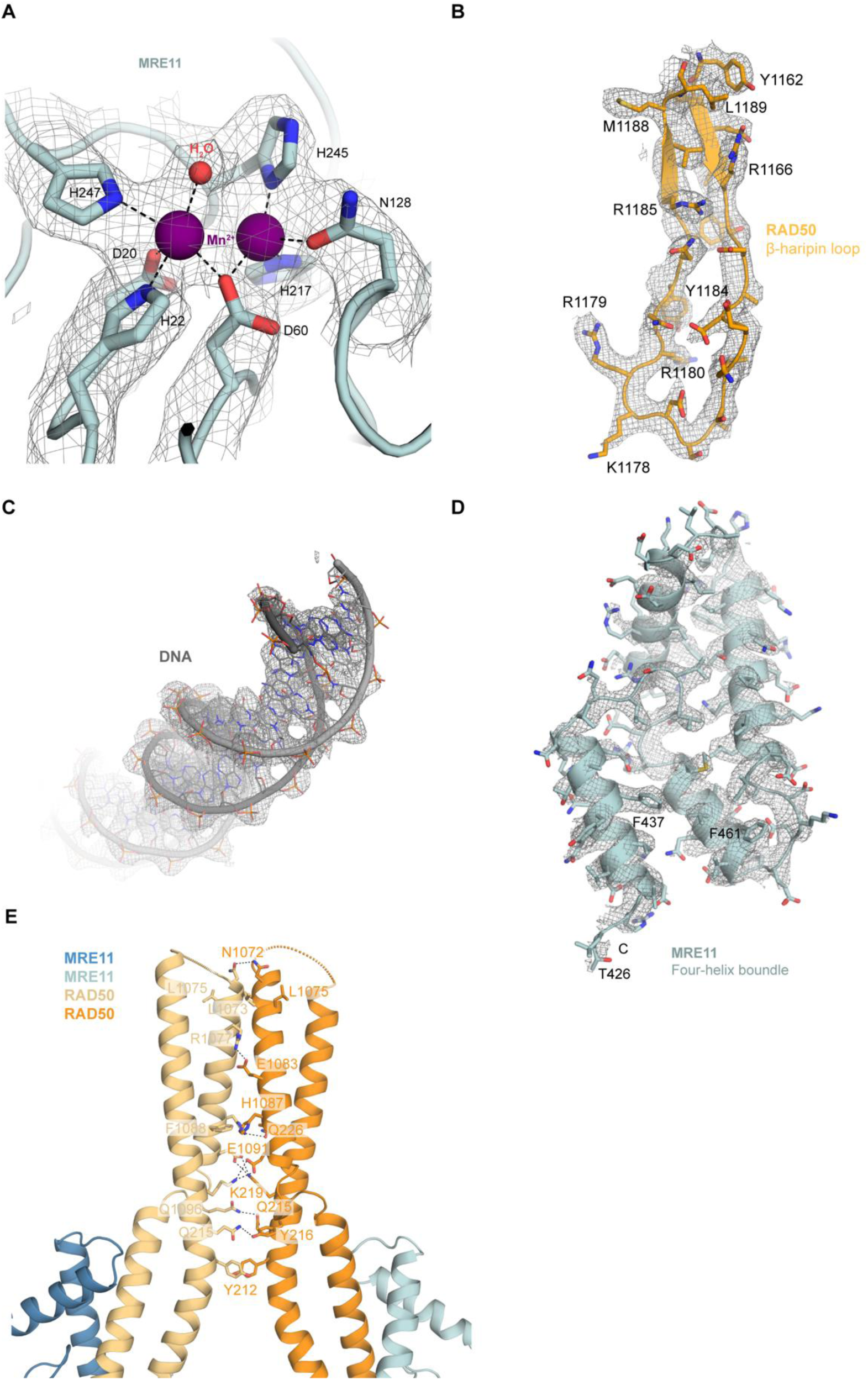
Quality of cryo-EM reconstructions. **A-E,** Densities for various parts of MR-TRF2^iDDR-Myb^-DNA complex as indicated. Interface of RAD50 CCDs with interacting residues shown as sticks and labeled.

**Supplementary Figure S5.**
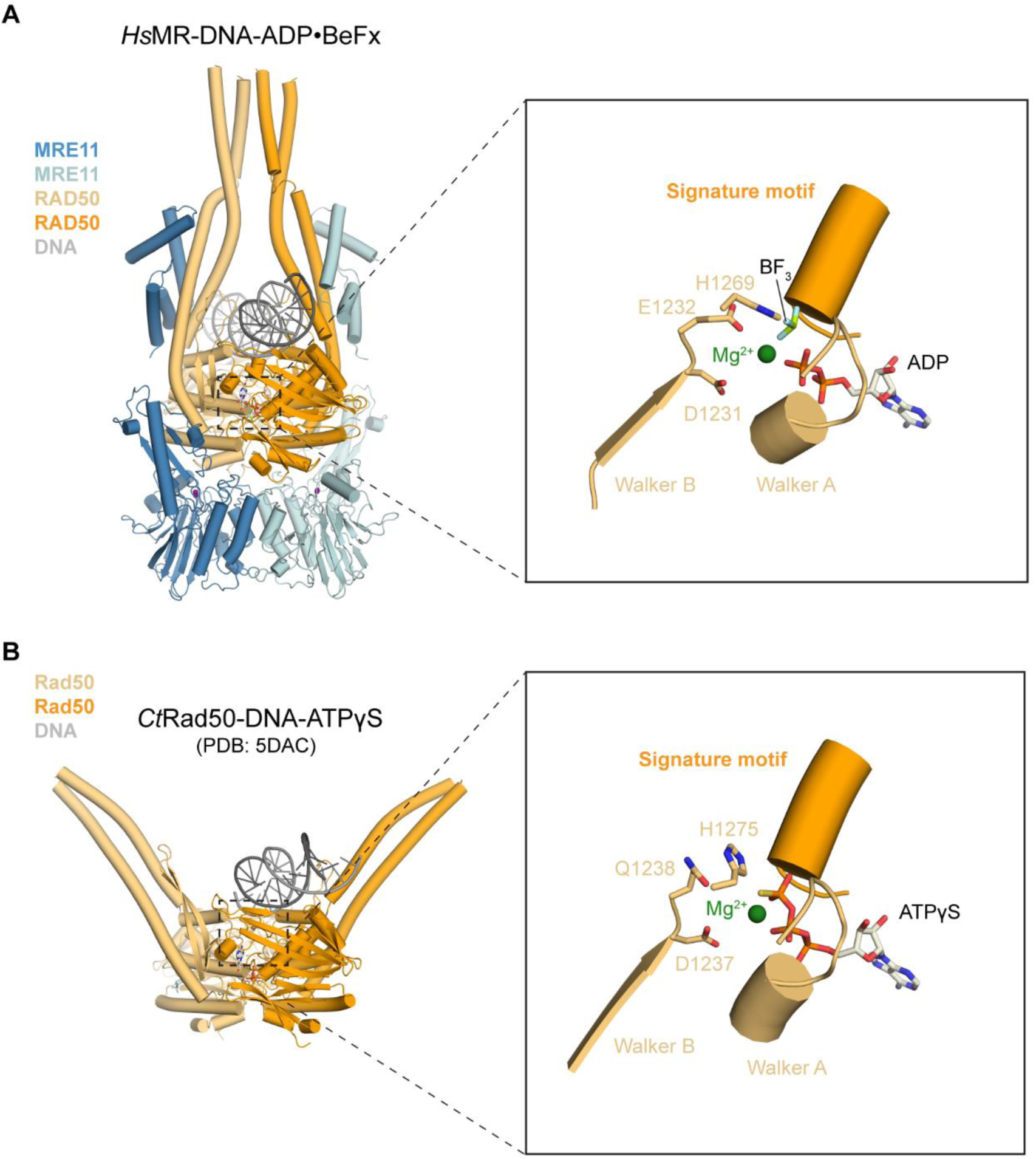
Structural comparison of nucleotide binding pockets in different (M)R structures. Side-by-side comparison of (M)R structures, emphasizing the unique structural arrangement of the signature motif surrounding RAD50’s nucleotide-binding site in **A,** *Hs*MR-DNA-ADP•BeF_x_ (current study) and **B,** *Ct*Rad50-DNA-ATPγS (PDB: 5DAC). Identically named subunits were colored consistently. *Hs*: *Homo sapiens, Ct*: *Chaetomium thermophilum*.

**Supplementary Figure S6.**
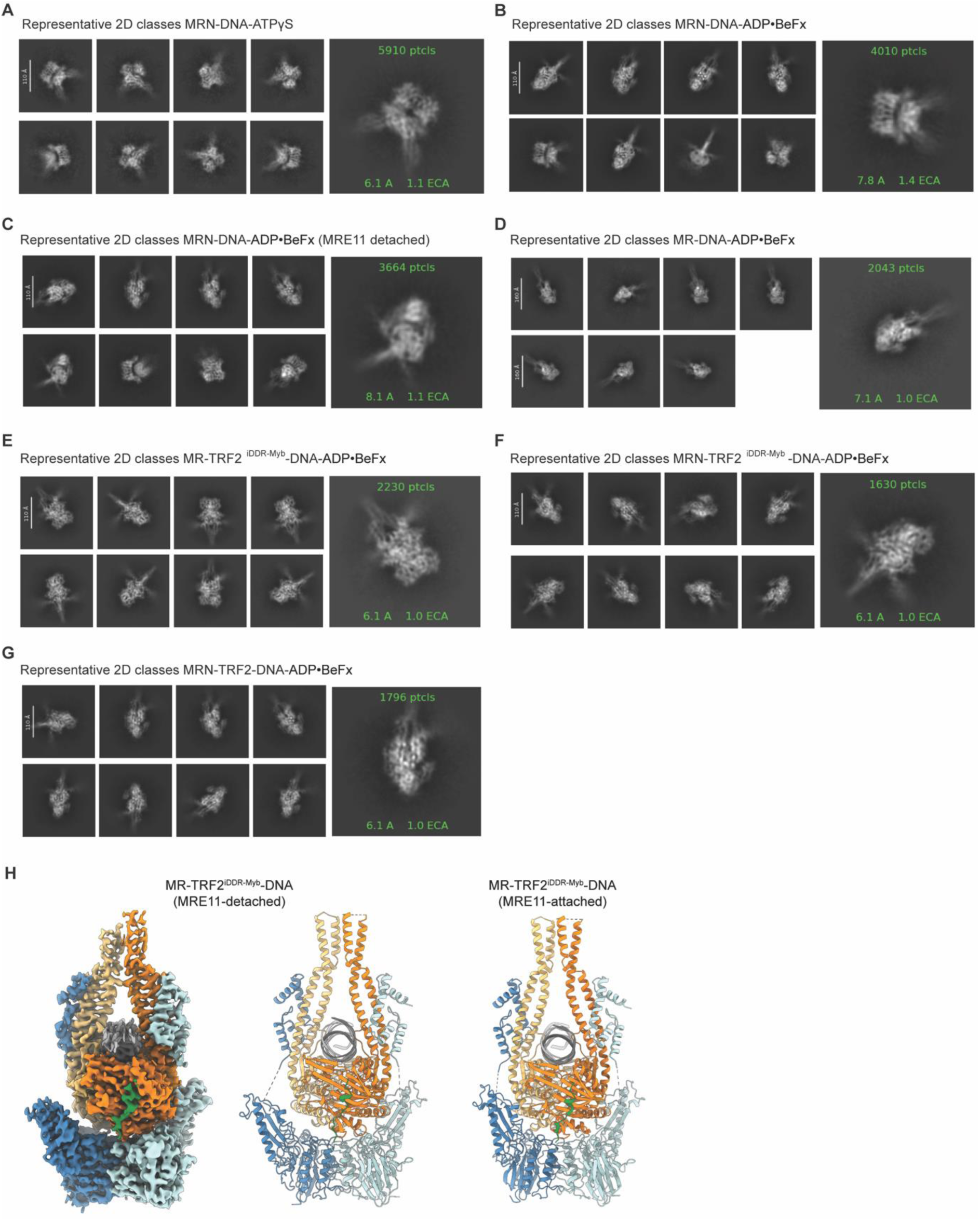
MR/MRN 2D classes. **A,** Representative 2D classes obtained for MRN with DNA and ATPγS. RAD50 coiled-coils domains (CCDs) are open. **B,** Representative 2D classes obtained for MRN with DNA and ADP•BeF_x._ The complex displayed a heterogeneity with open and closed CCDs, as well as asymmetrically bound MRE11 dimers. **C-G,** Representative 2D classes of MRE11 dimer detached as a rigid body from one RAD50^NBD^. **H,** Overview of the MR-TRF2iDDR-Myb-DNA complex structure with one partially detached MRE11 subunit (left two panels) and the side-by-side comparison to the fully attached MRE11 complex structure (rightmost panel). High resolution cryo-EM map of the partially detached MRE11 complex structure was displayed on the leftmost panel.

**Supplementary Figure S7:**
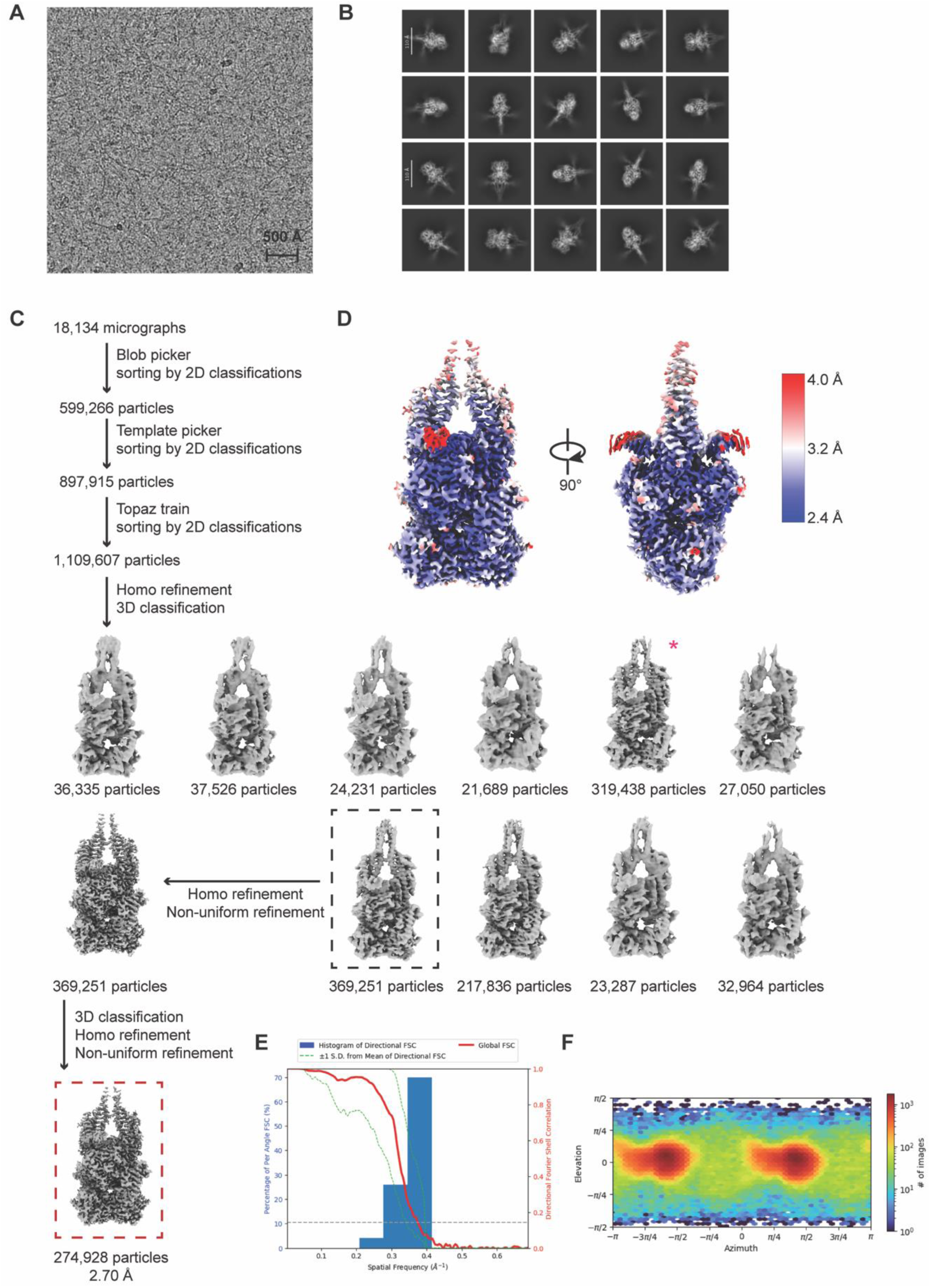
Cryo-EM data analysis of MR-TRF2^iDDR-Myb^-DNA. **A,** Representative micrograph of MR-TRF2^iDDR-Myb^-DNA among 18,134 collected movies. **B,** Representative 2D classes of the particles used for the final MR-TRF2^iDDR-Myb^-DNA reconstruction. **C,** Cryo-EM data processing workflow of MR-TRF2^iDDR-Myb^-DNA using cryoSPARC^83^. Pink asterisk denotes a well-resolved class with MRE11 detached from RAD50^NBD^. **D,** Local resolution visualisation of MR-TRF2^iDDR-Myb^-DNA calculated in cryoSPARC. Blue indicates higher resolution, red indicates lower resolution. **E,** Histogram of directional Fourier shell correlation (FSC)^91^ (blue) and global FSC curve (red) of the final MR-TRF2^iDDR-Myb^-DNA reconstruction. The spread of directional resolution values (green dashed lines) is defined as ± 1σ. The grey dashed line shows the 0.143 cut-off criterion, indicating a nominal resolution of 2.7Å. **F,** Angular distribution of the particles used for final MR-TRF2^iDDR-Myb^-DNA reconstruction.

**Supplementary Figure S8:**
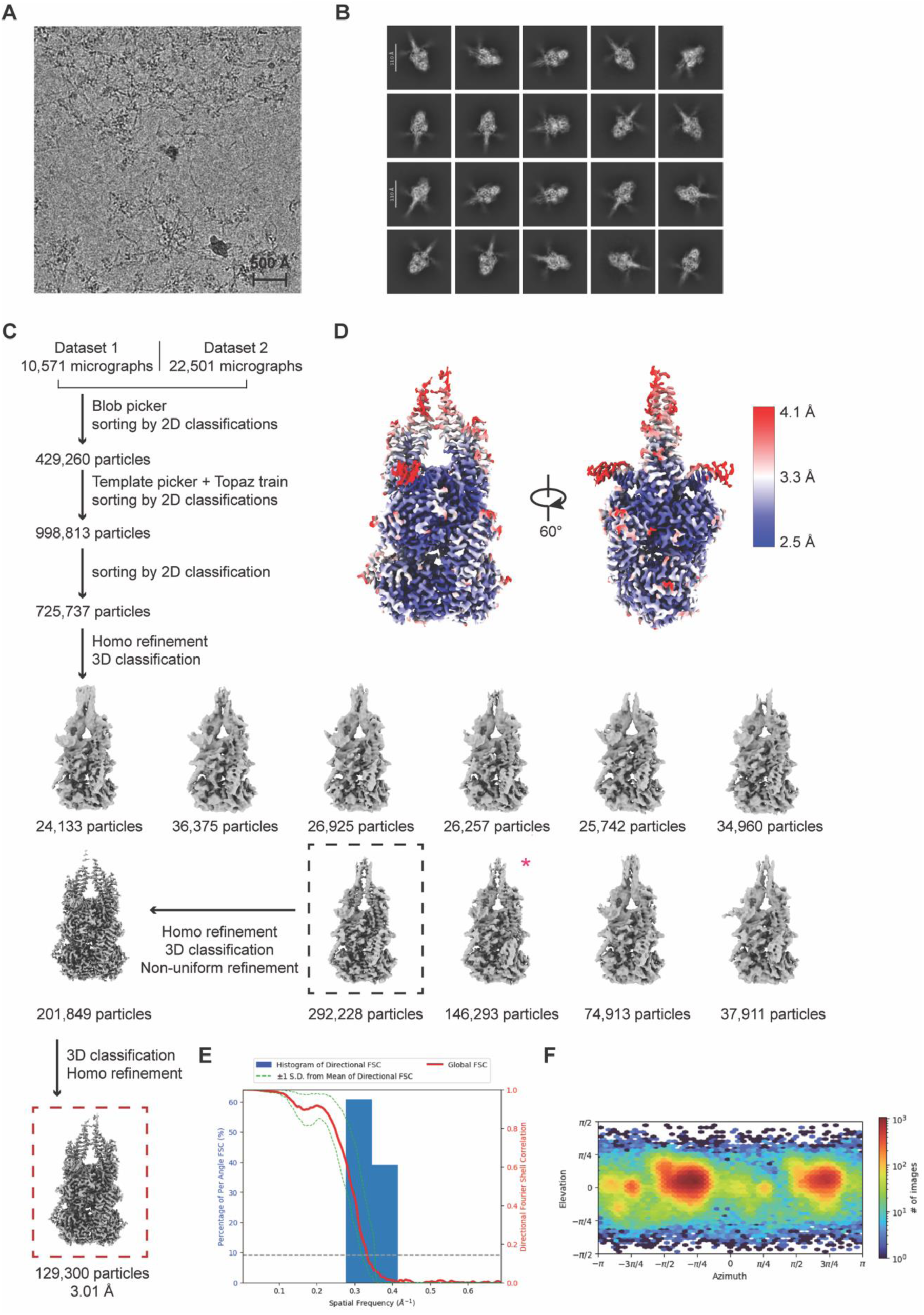
Cryo-EM data analysis of MRN-TRF2^iDDR-Myb^-DNA. **A,** Representative micrograph of MRN-TRF2^iDDR-Myb^-DNA among 33,072 collected movies. **B,** Representative 2D classes of the particles used for the final MRN-TRF2^iDDR-Myb^-DNA reconstruction. **C,** Cryo-EM data processing workflow of MRN-TRF2^iDDR-Myb^-DNA using cryoSPARC^83^. Pink asterisk denotes a well-resolved class with MRE11 detached from RAD50^NBD^. **D,** Local resolution visualisation of MRN-TRF2^iDDR-Myb^-DNA calculated in cryoSPARC. Blue indicates higher resolution, red indicates lower resolution. **E,** Histogram of directional Fourier shell correlation (FSC)^91^ (blue) and global FSC curve (red) of the final MRN-TRF2^iDDR-Myb^-DNA reconstruction. The spread of directional resolution values (green dashed lines) is defined as ± 1σ. The grey dashed line shows the 0.143 cut-off criterion, indicating a nominal resolution of 3.01Å. **F,** Angular distribution of the particles used for final MRN-TRF2^iDDR-Myb^-DNA reconstruction.

**Supplementary Figure S9:**
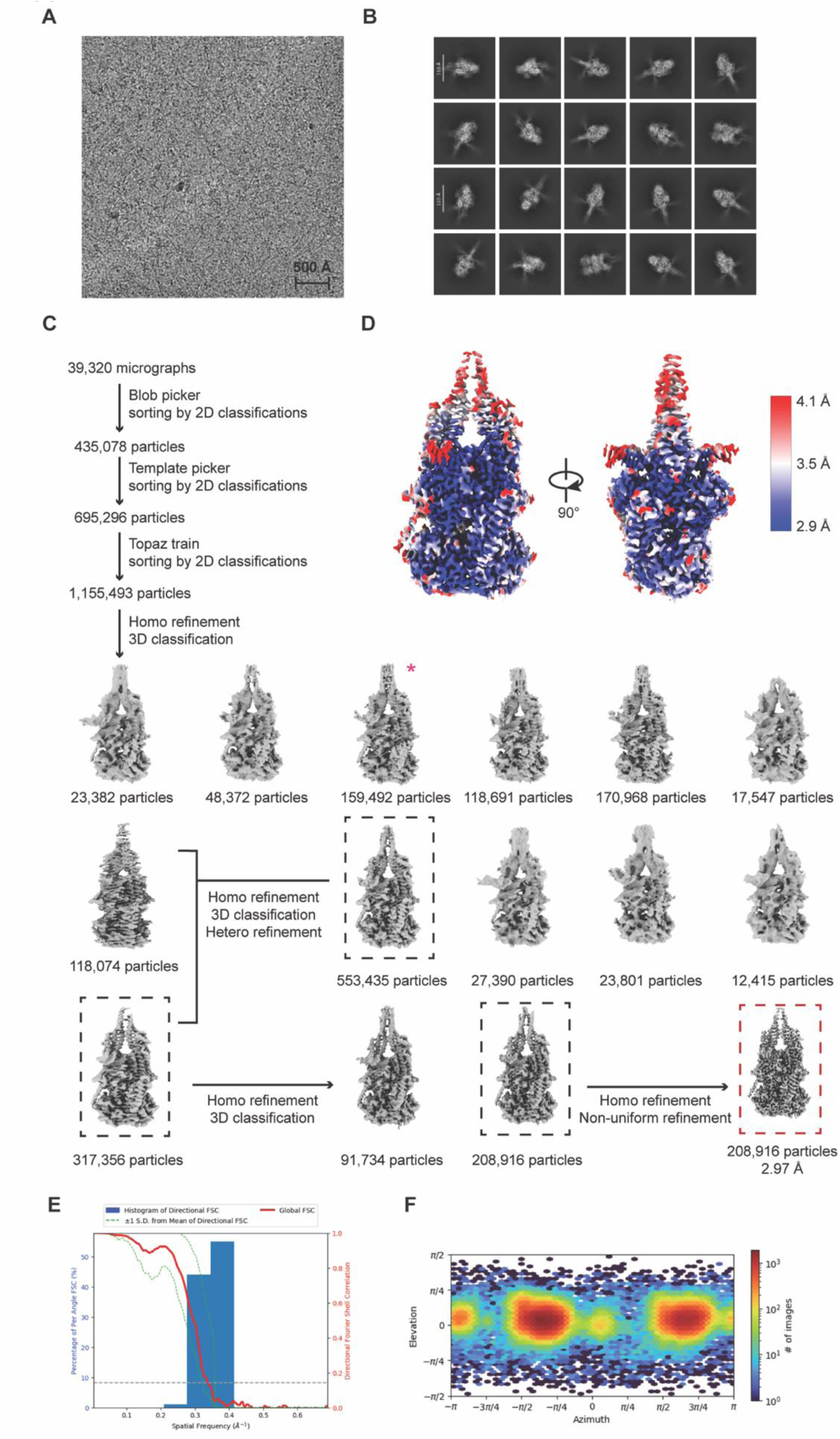
Cryo-EM data analysis of MRN-TRF2-DNA. **A,** Representative micrograph of MRN-TRF2-DNA among 39,320 collected movies. **B,** Representative 2D classes of the particles used for the final MRN-TRF2-DNA reconstruction. **C,** Cryo-EM data processing workflow of MRN-TRF2-DNA using cryoSPARC^83^. Pink asterisk denotes a well-resolved class with MRE11 detached from RAD50^NBD^. **D,** Local resolution visualisation of MRN-TRF2-DNA calculated in cryoSPARC. Blue indicates higher resolution, red indicates lower resolution. **E,** Histogram of directional Fourier shell correlation (FSC)^91^ (blue) and global FSC curve (red) of the final MRN-TRF2-DNA reconstruction. The spread of directional resolution values (green dashed lines) is defined as ± 1σ. The grey dashed line shows the 0.143 cut-off criterion, indicating a nominal resolution of 2.97Å. **F,** Angular distribution of the particles used for final MRN-TRF2-DNA reconstruction.

**Supplementary Figure S10:**
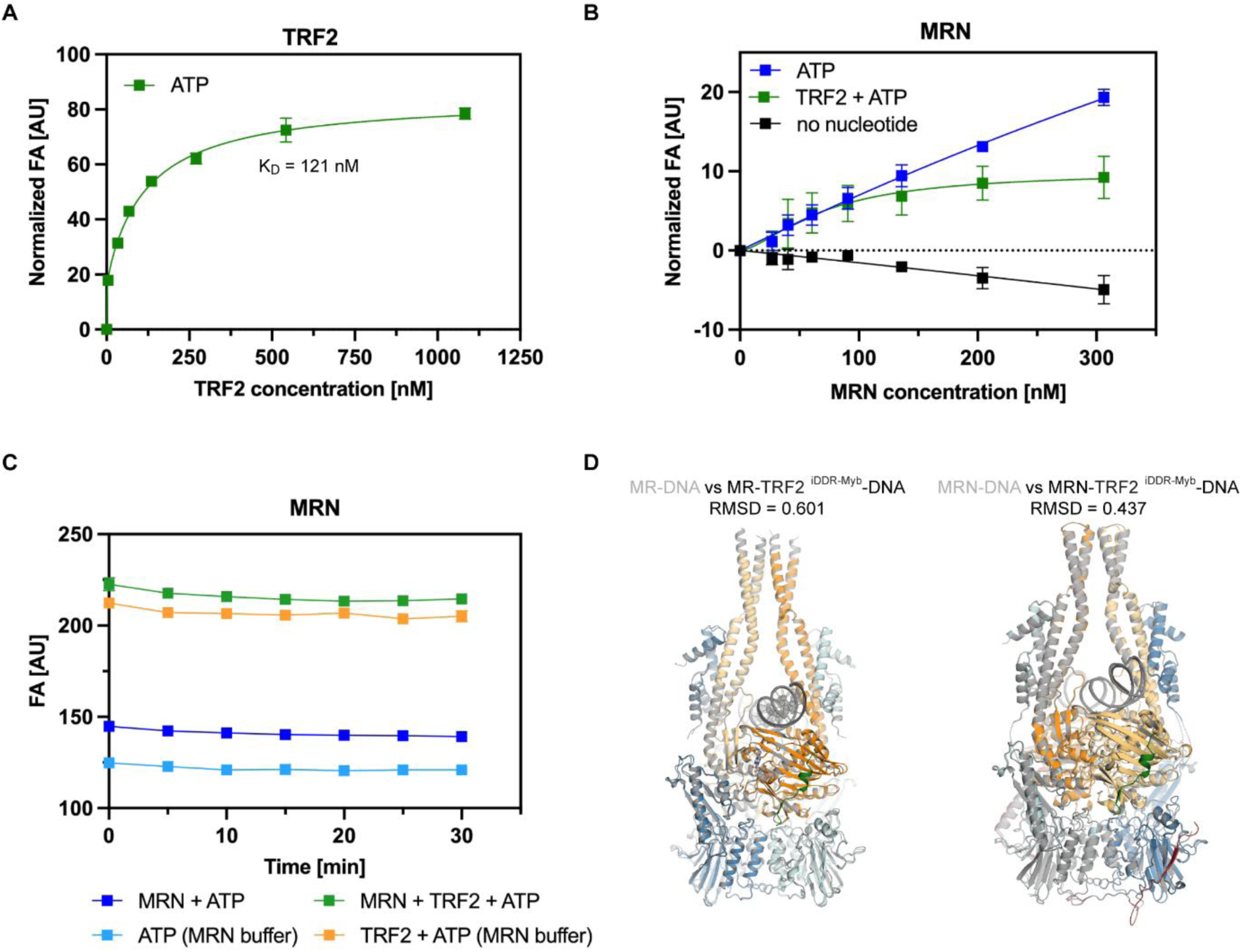
Combined results of binding assays and structural comparison of MR(N)-DNA complex with MR(N)-TRF2^iDDR-Myb^-DNA complex. **A,** Fluorescence anisotropy (FA)-based assay assessing full length TRF2 binding to 30bp-Myb DNA in the presence of ATP. The experiment was carried out in three independent replicates and data are represented as mean values ± SD. K_D_ value was derived by fitting the anisotropy data to two sites specific binding with Hill slope model in Prism (GraphPad). **B**, FA-based assay documenting the effects of ATP, as well as ATP and TRF2 on MRN binding towards telomeric repeat containing 64-mer DNA template. Each experiment was performed in three independent replicates and data are represented as mean values ± SD. **C**, FA signals of each titration condition kept relatively stable over the measurement time period. A concentration of 300 nM MRN and/or 1.6 μM TRF2 were used where indicated. **D**, Structural superimposition of MR(N)-DNA complex with MR(N)-TRF2^iDDR-Myb^-DNA complex. MR(N)-DNA complex structure was colored in grey for simplicity, while MR(N)-TRF2^iDDR-Myb^-DNA complex structure was colored consistently as other figures in the manuscript.

**Supplementary Table S1.**

|  | MR-DNA<br>(EMDB-52959)<br>(PDB 9Q9H) | MRN-DNA<br>(EMDB-52960)<br>(PDB 9Q9I) | MR-TRF2<sup>iDDR</sup>-<br>Myb-DNA<br>(EMDB-52962)<br>(PDB<br>9Q9K) | MRN-TRF2<sup>iDDR</sup>-<br>Myb-DNA<br>(EMDB-52961)<br>(PDB<br>9Q9J) | MRN-TRF2-DNA<br>(EMDB-52964)<br>(PDB<br>9Q9M) |
| --- | --- | --- | --- | --- | --- |
| **Data collection and processing** |  |  |  |  |  |
| Magnification | 130,000 | 165,000 | 165,000 | 165,000 | 165,000 |
| Voltage (kV) | 300 | 300 | 300 | 300 | 300 |
| Electron exposure (e-/Å<sup>2</sup>) | 42.377 | 40 | 40 | 40 | 40 |
| Defocus range (μm) | -1.1 to -2.9 | -0.5 to -2.6 | -0.5 to -2.6 | -0.5 to -2.6 | -0.5 to -2.6 |
| Pixel size (Å) | 1.06 | 0.727 | 0.727 | 0.727 | 0.727 |
| Symmetry imposed | C1 | C1 | C1 | C1 | C1 |
| Initial particle images (no.) | 817,041 | 723,315 | 1,109,607 | 725,737 | 1,155,493 |
| Final particle images (no.) | 95,397 | 172,784 | 274,928 | 129,300 | 208,916 |
| Map resolution (Å) | 3.21 | 3.08 | 2.70 | 3.01 | 2.97 |
| FSC threshold | 0.143 | 0.143 | 0.143 | 0.143 | 0.143 |
| **Refinement** |  |  |  |  |  |
| Initial model used (PDB code) | AlphaFold3 | AlphaFold3 | 9Q9H | 9Q9I | 9Q9J |
| Model resolution (Å) | 3.2 | 3.2 | 2.9 | 3.1 | 3.2 |
| FSC threshold | 0.5 | 0.5 | 0.5 | 0.5 | 0.5 |
| Map sharpening *B* factor (Å<sup>2</sup>) | -110 | -85 | -89 | -87 | -80 |
| **Model composition** |  |  |  |  |  |
| Non-hydrogen atoms | 16,786 | 17,121 | 16,910 | 17,465 | 17,361 |
| Protein residues | 1,932 | 1,960 | 1,934 | 2,004 | 1,993 |
| Nucleotides | 50 | 54 | 54 | 52 | 52 |
| ***B* factors (Å<sup>2</sup>)** |  |  |  |  |  |
| Protein | 62.66 | 110.93 | 60.01 | 78.74 | 78.55 |
| Nucleotide | 99.03 | 93.51 | 46.38 | 52.12 | 52.15 |
| Ligand | 24.04 | 92.97 | 34.21 | 54.76 | 54.76 |
| Water | 24.41 | 82.39 | 28.03 | 44.38 | 44.38 |
| **R.m.s. deviations** |  |  |  |  |  |
| Bond lengths (Å) | 0.006 | 0.006 | 0.006 | 0.006 | 0.006 |
| Bond angles (°) | 1.203 | 1.161 | 1.161 | 1.076 | 1.174 |
| **Validation** |  |  |  |  |  |
| MolProbity score | 2.09 | 2.02 | 1.93 | 1.71 | 1.95 |
| Clashscore | 10.77 | 10.65 | 10.48 | 8.18 | 10.22 |
| Poor rotamers (%) | 0.8 | 0.7 | 0.5 | 0.4 | 0.5 |
| **Ramachandran plot** |  |  |  |  |  |
| Favored (%) | 95.77 | 95.92 | 96.81 | 97.53 | 96.65 |
| Allowed (%) | 4.23 | 4.08 | 3.19 | 2.47 | 3.35 |
| Disallowed (%) | 0.00 | 0.00 | 0.00 | 0.00 | 0.00 |

